# Adaptation of protein stability to thermally heterogeneous environments

**DOI:** 10.1101/2025.02.12.637818

**Authors:** Tadeas Priklopil, Kirsten Bomblies, Alex Widmer

## Abstract

Proper protein folding is essential for biological function, and its disruption can lead to disease, reduced fitness, or death. The ability of a protein to maintain its folded conformation is thus critical for life, making it a key target of adaptive evolution. However, protein stability is sensitive to environmental factors, particularly temperature, which can threaten phenotypic integrity and organismal survival under thermal changes. Despite its importance, the influence of complex thermal environments – characterized here by mean temperature, thermal fluctuations, and environmental heterogeneity – on the evolution of protein stability remains poorly understood. To address this, we developed a mathematical framework that combines two well-established models: a population genetic model describing species distributed across habitats with distinct thermal environments, and a thermodynamic model of protein stability incorporating temperature-dependent enthalpy and entropy contributions. We focus on two-state proteins that alternate between folded and unfolded states and assume that allelic fitness is maximized in proteins that achieve an optimal balance between flexibility and rigidity. Using this framework, we performed an invasion analysis of mutations (*sensu* adaptive dynamics framework) affecting three thermodynamic parameters that fully determine protein stability profiles. Where possible, we derived analytical expressions for evolutionarily optimal thermodynamic parameters and complemented these with numerical solutions. Our results show that mean temperature and thermal fluctuations have orthogonal effects on thermodynamic parameters, underscoring the need to consider both when studying protein stability adaptation. We further examined thermally heterogeneous environments, where subpopulations connected by migration experience different mean temperatures, identifying conditions that favor either local (specialist) or global (generalist) adaptation. Our results may explain why one thermodynamic parameter shows little association with thermal adaptation and suggest that local adaptation is more likely for proteins with stability profiles limited to narrow temperature ranges. Additionally, our analysis reveals whether a locally adapted protein originated in a colder or warmer habitat. Finally, we identified trade-offs in thermodynamic parameters that influence local or global adaptation. This study offers key predictions about protein evolution in complex thermal environments and lays the groundwork for developing practical tools to understand how temperature shapes adaptation and biodiversity.

## 1 Introduction

Temperature is a fundamental driver of evolutionary processes, shaping the physiological and biochemical traits of organisms across diverse habitats (Angilletta, 2009). Over evolutionary time, species have adapted to a wide spectrum of thermal conditions, from the extreme cold of Arctic regions to the searing heat of volcanic environments (Somero et al., 2017). However, despite this remarkable adaptability, most species remain confined to relatively narrow thermal ranges and can be vulnerable to even slight deviations from their preferred temperatures (Oliver and Palumbi, 2011; Somero et al., 2017). Understanding the mechanisms of thermal adaptation is therefore critical, particularly as rapid environmental changes impose new thermal challenges on natural populations.

Proteins, which are fundamental to nearly every cellular function, are highly sensitive to changes in temperature. These molecules facilitate a broad range of biological processes, including enzymatic catalysis and molecular signaling, all of which rely on maintaining their specific, stable three-dimensional structures (Dill et al., 2017; Alberts et al., 2022). However, maintaining protein stability presents a challenge in changing thermal environments. Proteins must balance flexibility, which allows for conformational changes necessary for function, and rigidity, which ensures structural integrity (Somero et al., 2017). Temperature shifts can upset this balance, possibly leading to denaturation resulting in impaired cellular processes or even organismal death (Fersht, 1999; Dill et al., 2017).

The ability of proteins to remain stable under varying thermal conditions is a key factor in the thermal adaptation of organisms (DePristo et al., 2005; Sikosek and Chan, 2014; Echave and Wilke, 2017; Bastolla et al., 2017; Bershtein et al., 2017). Although various cellular mechanisms, such as chaperone activity or heat shock responses, can assist in maintaining protein stability under thermal stress (Alberts et al., 2022), protein stability itself can evolve in response to shifts in thermal environments (Zavodszky et al., 1998; Hochachka and Somero, 2002; Fields et al., 2015; Barik, 2020). Indeed, several studies demonstrate that genetic differentiation related to protein stability often correlates with habitat temperature. For example, the cytosolic malate dehydrogenase (cMDH) enzyme, a key enzyme in the Krebs cycle, exhibits temperature-associated polymorphism across multiple genera of marine molluscs (Dong et al., 2018; Liao et al., 2019), as well as in honeybees (Meemongkolkiat et al., 2020) and the plant *Arabidopsis thaliana* (Simon et al., 1983). In the honeybee and plant examples, alternate alleles are adaptive to different latitudes, and the encoded proteins display temperature-dependent differences in activity. Similarly, polymorphisms affecting the stability of the phosphoglucose isomerase (Pgi) enzyme, associated with temperature, have been observed in the Glanville fritillary butterfly (Kallioniemi and Hanski, 2011; De Jong and Saastamoinen, 2018; Yang et al., 2023). In bacteria, protein stability effectively describes the thermal responses of 35 bacterial species (Chen and Shakhnovich, 2010), and recent in vitro selection experiments have documented mutations that improve protein stability as a cold-adapted bacterium is subjected to gradually increasing temperatures (Toll-Riera et al., 2022). Remarkably, even a single amino acid substitution affecting protein stability can correlate with thermal environments (Dong and Somero, 2009).

Much research has focused on large thermal differences between environments (see also reviews by Jaenicke, 1991; Fields, 2001; Feller, 2018), but recent evidence suggest that even modest changes in temperature can produce genetic variation in protein stability. For instance, in warm- and cold-adapted marine mussel species, a large number of proteins assessed for stability-temperature correlation demonstrated that a difference as small as 4°C in the critical temperature of the organism can lead to stability-associated genetic variation (Chao et al., 2020). However, despite these insights, empirical data on how small temperature differences drive thermal adaptation remain limited.

Thermal adaptation is further complicated by the fact that populations subjected to similar thermal stresses might not be geographically isolated, leading to gene flow between populations. Gene flow can weaken local adaptation by introducing alleles that are not suited to local thermal conditions, potentially hindering the action of natural selection (Kawecki and Ebert, 2004; Blanquart et al., 2013). Additionally, thermal fluctuations within an environment likely influence the course of thermal adaptation. For example, latitudinal variations in the relationship between thermal optima and mean habitat temperature (Deutsch et al., 2008; Tewksbury et al., 2008) have been linked to differences in the magnitude of thermal fluctuations (Amarasekare and Johnson, 2017). However, there is limited theoretical or empirical research into how these factors – mean temperature, thermal fluctuations, and gene flow – interact to influence the evolution of protein stability. Developing mathematical models to investigate these interactions could offer valuable insights, helping to bridge gaps in our understanding of thermal adaptation (see e.g., Berger et al., 2021). This would be especially important given the substantial genetic variation observed along thermal gradients (Angilletta, 2009), raising the key question of how much of this variation is due to biophysical adaptations in protein stability.

In this study, we address critical gaps in our understanding of the thermal adaptation of protein stability by developing a mathematical framework that integrates two well-established models. The first is a population genetic model describing a species distributed across multiple habitats with distinct thermal environments (Bulmer, 1972; Bürger, 2000), and the second is a model of protein thermodynamic stability that incorporates temperature-dependent enthalpy and entropy contributions (Becktel and Schellman, 1987; Barrick, 2018). This framework incorporates three key environmental variables: mean temperature, the magnitude of thermal fluctuations, and thermally heterogeneous, interconnected environments. It assumes that allelic fitness is maximized when a protein remains in its optimally folded state for a defined proportion of time between synthesis and degradation, ensuring a balance between rigidity and flexibility (Somero et al., 2017), a principle well-established in studies of protein evolution (Fields, 2001; DePristo et al., 2005; Williams et al., 2006; Shah et al., 2015; Somero et al., 2017; Echave and Wilke, 2017; Agozzino and Dill, 2018; Berger et al., 2021). Protein stability is defined in terms of thermodynamic stability, based on Gibbs free energy, which directly links temperature to protein stability. By focusing on two-state model proteins that alternate between folded and unfolded states, we capture the folding dynamics of many single-domain proteins (Zwanzig, 1997; Privalov, 2012; Dill et al., 2017; Barrick, 2018). This approach is practical because their stability temperature profiles can be fully described using only three thermodynamic parameters – maximum stability, the temperature of maximum stability, and heat capacity change – which are the focus of our evolutionary analysis. Additionally, these parameters can be estimated experimentally (Fersht, 1999; Barrick, 2018; Thurlkill et al., 2023) or computationally (Seeliger and De Groot, 2010; Kim et al., 2016; Timr et al., 2020; Galano-Frutos et al., 2023; Wang et al., 2024), allowing us to link genetic variation directly to adaptive phenotypes.

Our approach combines analytical and numerical solutions to provide new insights into the evolutionary dynamics of protein stability. We explore how mean temperature, thermal fluctuations, and thermal heterogeneity drive the adaptive evolution of the three key thermodynamic parameters. Our findings reveal that while mean temperature is the primary driver of adaptation, thermal fluctuations must also be considered, as they exert orthogonal effects on thermodynamic parameters. We observe local adaptation when thermal environments differ significantly between habitats, with stronger signals of adaptation emerging when populations adapt to warmer environments after evolving in colder ones, compared to the reverse scenario. In some cases, increasing thermal fluctuations further promotes local adaptation. We also identify trade-offs between thermodynamic parameters that shape the balance between local and global adaptation. Finally, we compare these findings to a simpler model in which enthalpy and entropy contributions are independent of temperature, revealing that temperature-dependence introduces significant complexity and leads to more intricate adaptive responses. Together, these results address critical gaps in our understanding of protein adaptation and provide broader insights into how organisms adapt to and cope with changing environmental conditions.

The paper is organised as follows: In Section 2, we introduce the population genetic modelling framework and the mechanistic derivation of protein stability. In Section 3, we provide a general outline of the mathematical methods to analyse the evolutionary dynamics of protein stability and then present the results. In Section 3.1, we present the evolutionarily favoured thermodynamic parameters in a thermally fluctuating but homogeneous (“single habitat”) environment. In Section 3.2, we investigate the conditions under which local adaptation is favoured in a thermally heterogeneous (“two habitat”) environment. After presenting the general results, we focus on two typical biological scenarios: one where a warmer habitat becomes available to an organism originally adapted to a colder habitat, and the other where a colder habitat becomes available to an organism adapted to a warmer habitat. In Section 3.2.4, we investigate the role of thermodynamic trade-offs in the adaptive process. For both homogeneous and heterogeneous environments, we compare our results to a simpler model where enthalpy and entropy contributions are independent of temperature (Sections 3.1.3 and 3.2.5). All results are presented in terms of thermodynamic, demographic, and environmental parameters. Further details on the mathematical model and its analysis can be found in the Appendix and the Supplementary material.

## 2 Model

Consider a large, constant population subdivided into a finite number of habitats A, B, Individuals are assumed to be haploid, with non-overlapping discrete generations, and follow a life cycle consisting of viability selection, reproduction, population regulation, and migration.

Viability selection occurs at the start of each generation and spans from birth to reproduction. During this phase, individuals experience habitat-specific thermal fluctuations of magnitude *ϵ*_A_, *ϵ*_B_,… around habitat-specific average temperatures 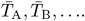 Selection depends on the biophysical and thermodynamic properties of the protein of interest, which are fully determined by a set of thermodynamic parameters collectively denoted as ***ϕ***. These parameters, in turn, are determined by the alleles at a single locus under consideration. Within each habitat, viability selection is assumed to depend on the difference between the temperature-dependent thermodynamic stability Δ*G* of the protein and a temperature-dependent optimal protein stability Δ*G*^opt^, which maximizes the protein’s functional role (further details on protein stability are provided in Section 2.1). This difference directly determines the performance of the phenotype of interest and, consequently, the probability of survival during viability selection. The probability that offspring survive viability selection is

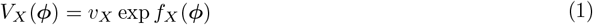

in habitat *X* ∈ {A, B,}, where *v*_*X*_ is the maximum surviving probabilities in the habitat, and where

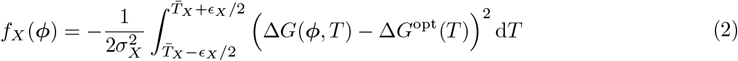

gives a scaled distance between the stability profile of the protein of interest Δ*G*(***ϕ***) and its optimal stability profile Δ*G*^opt^ over the thermal fluctuations in habitat *X* ∈ {A, B, …}. Whenever we consider a model with small (negligible) thermal fluctuations, we use 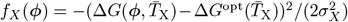.

The scaling parameter *σ*_*X*_ controls how sensitive viability selection is to protein stability in habitat *X* ∈ {A, B, …}, which in this paper is assumed to be equal across all habitats *σ* = *σ*_*X*_ for all *X* ∈ {A, B, …}. The probability of surviving viability selection (eq. 1) thus increases with increasing *σ*, and ‘closer’ the stability profile of the protein Δ*G*(***ϕ***) is to its optimal stability profile Δ*G*^opt^ in each thermal environment (Figure 1).

**Figure 1:**
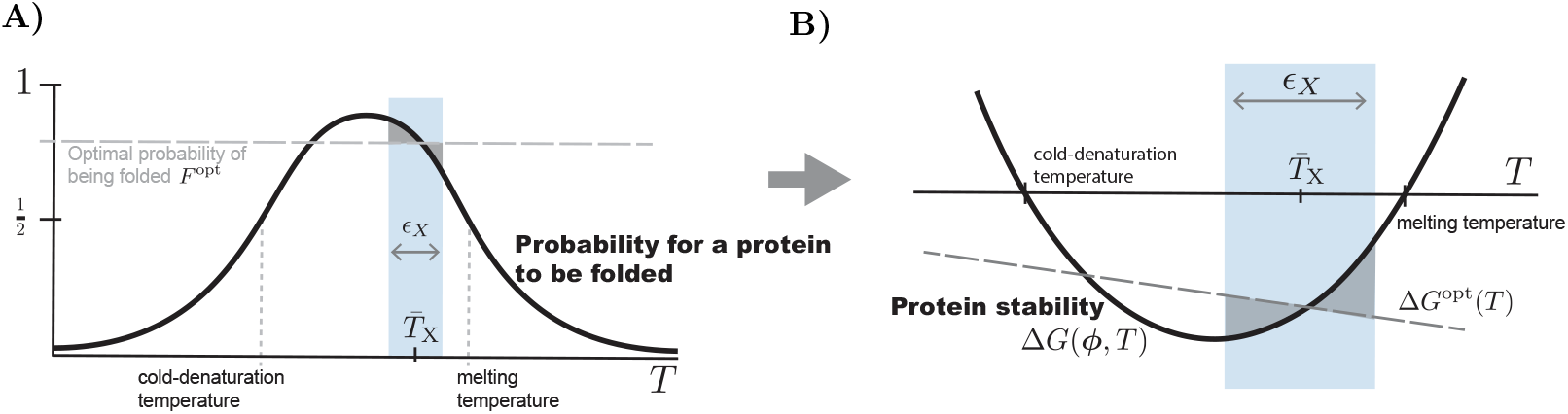
**A)** Probability for a two-state protein to be in its folded state as a function of temperature *T*. Performance of the phenotype is optimal when the probability is *F* ^opt^ (0 *< F* ^opt^ *<* 1). With blue color we have indicated the range of temperatures 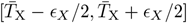 the organism experiences in habitat *X*. The shaded areas indicate the difference between the optimal probability and the probability to be folded over the range of temperatures. Note that this bell-shaped curve has two denaturation temperatures where exactly half of the proteins are folded, one for higher temperatures (heat denaturation or melting) and the other for lower temperatures (cold denaturation). This panel is translated into the thermodynamical description of protein stability in panel B. **B)** Protein stability profile Δ*G*(***ϕ***) of a two-state protein, depicted with a thick black curve, is a convex function of temperature *T* (its dependence on thermodynamical parameters ***ϕ*** is explored in Figure 2). The more stable the protein is at *T*, the more negative the value of Δ*G*(***ϕ***, *T*). The decreasing straight dashed line depicts the optimal protein stability Δ*G*^opt^(*T*) = − *RT* ln *K*^opt^ where *K*^opt^ = *F* ^opt^*/*(1 − *F* ^opt^) (eq. 7 in Section 2.1) and corresponds to the horizontal line at *F* ^opt^ in panel A. The color-sheme is identical to panel A, with gray regions indicating how close is protein stability to its optimal stability over the thermal fluctuations in habitat *X*, with the smaller area indicating a smaller distance and hence greater survival probability (eqs. 1-2).

After viability selection, all surviving offspring mature into reproducing adults, each of which produces a large number of offspring and then dies. The offspring then undergo non-selective competition to maintain the habitat-specific population size, and then either stay in their native habitat or disperse to another habitat. The probability that an offspring in habitat *X* ∈ {A, B, …} originated from habitat *Y* ∈ {A, B, …} is denoted by *m*_*XY*_, which we assume to be independent of protein stability. After dispersal, a new generation begins. Supposing that alleles *a, b*, … segregate across the populations with frequencies *p*_*X,a*_, *p*_*X,b*_, …, with 1 = ∑_*x*∈*{a,b*,…}_ *p*_*X,i*_ in each habitat *X* ∈ {A, B, …}, the population genetics of allele *x* ∈ *{a, b*, …} follows the recursion

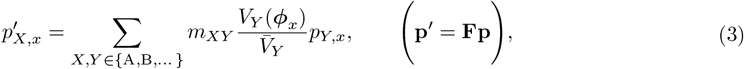

where 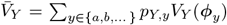 is the average probability of surviving in habitat *Y* ∈ {A, B, …}, and where *m*_*XX*_ = 1 ∑− _*Y* ∈{A,B,…}_ *m*_*XY*_ for all *X* ∈ {A, B, …} (Appendix A). The equation in the brackets expresses the equations in a matrix notation, with each row given by the equality on the left. Note that we have used the notation ***ϕ***_*x*_ to indicate that thermodynamical parameters depend on the genetic sequence *x* of the allele at the locus of interest. Also note that in eq. (3), frequency 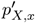 can be interpreted as the probability that a randomly sampled allele in *X* in this generation came from *Y* in the previous generation and is an *x* that survived viability selection, 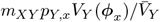,and then summed over all possible habitats of origin (right-hand-side of eq. 3). The migration model (eq. 3) is a version of models analysed in Bulmer, 1972; Christiansen and Feldman, 1975; Nagylaki and Lou, 2008; Yeaman and Otto, 2011.

The central role in our mathematical analysis is invasion fitness, which for the allele *x* ∈ {*a, b*, …} is the dominant eigenvalue of the matrix **F** that is linearized about an equilibrium where the allele *x* is absent (we assume the model contains an equilibrium at which the remaining alleles can stably coexist). Whenever invasion fitness is larger than 1, the allele *x* can invade into the population, otherwise it can not (Tuljapurkar, 1989; Metz et al., 1992).

### 2.1 Protein stability and thermodynamical parameters

Here, we derive a representation of the protein stability temperature profile, Δ*G*(***ϕ***), which depends on thermodynamic parameters collected in the vector ***ϕ***. To derive Δ*G*(***ϕ***), we assume that our protein of interest is a two-state protein. This means that, at any point in time from synthesis to degradation, it is either in the folded state F or the unfolded state U, and can transition between these states (Zwanzig, 1997). Most small and single-domain proteins are well described by this two-state model (Privalov, 2012). The thermodynamic stability of such two-state proteins can be defined as the difference in Gibbs free energy, Δ*G*(***ϕ***, *T*) = *G*_F_(***ϕ***, *T*) − *G*_U_(***ϕ***, *T*), between its folded F and unfolded U states at temperature *T* (Privalov and Khechinashvili, 1974; Fersht, 1999; Barrick, 2018). Protein stability at *T* is composed of enthalpic and entropic contributions as

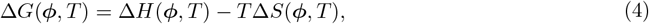

where Δ*H*(***ϕ***, *T*) = *H*_F_(***ϕ***, *T*) − *H*_U_(***ϕ***, *T*) and Δ*S*(***ϕ***, *T*) = *S*_F_(***ϕ***, *T*) − *S*_U_(***ϕ***, *T*) represent the differences in enthalpy and entropy, respectively, between folded and unfolded proteins at temperature *T*. The enthalpic contribution primarily reflects the strength of stabilizing interactions such as hydrogen bonds, van der Waals forces, and ionic bonds within the folded protein (Fersht, 1999; Dill et al., 2017; Barrick, 2018). The entropic contribution, on the other hand, arises from the disorder in the system, with unfolded proteins having higher entropy due to their flexible, random conformations. Key determinants of these contributions include hydrophobic residues, which promote folding by minimizing water contact and increasing enthalpic stability, and flexible residues like glycine and proline, which influence entropy by affecting protein backbone flexibility (Fersht, 1999; Dill et al., 2017; Barrick, 2018; Hait et al., 2020).

Enthalpic and entropic contributions in eq. (4) depend on temperature *T*, and can be expressed in terms of the heat capacity of the protein (Barrick, 2018). For a protein in state F, the expression (∂*H*_F_*/*∂*T*)_p_ = *C*_p,F_ relates heat capacity and enthalphy, and *T* (∂*S*_F_*/*∂*T*)_p_ = *C*_p,F_ relates heat capacity and entropy, and where the subscript p indicates that these are defined under constant pressure. Similar expressions hold for a protein in state U. We assume that Δ*C*_p_ = *C*_p,F_ −*C*_p,U_ is to a good approximation independent of temperature (Becktel and Schellman, 1987; Schellman, 1987; McCrary et al., 1996; Barrick, 2018). This then leads to protein stability (eq. 4) that is fully determined by three thermodynamical parameters ***ϕ*** = (Δ*G*_max_, Δ*C*_p_, *T*_max_), and that to a good approximation can be expressed as

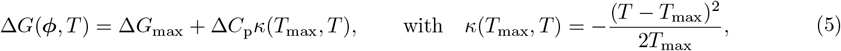

where the parameter Δ*G*_max_ := Δ*H*(*T*_max_) is the difference in enthalpy evaluated at *T*_max_ (Appendix A.2). Equation (5) yields a protein stability curve that forms an upward parabola whenever Δ*C*_p_ *<* 0, a condition we assume is always satisfied, and it attains its minimum Δ*G*_max_ *<* 0 at the reference temperature *T*_max_ (Figure 2). We therefore refer to the thermodynamic parameter Δ*G*_max_ as the maximum stability and *T*_max_ as the maximum-stability temperature. Furthermore, the thermodynamic parameter representing the heat capacity change between folded and unfolded proteins, Δ*C*_p_, scales the temperature dependence of protein stability (eq. 5) and thus determines the width of the stability curve. We refer to Δ*C*_p_ simply as the heat capacity.

**Figure 2:**
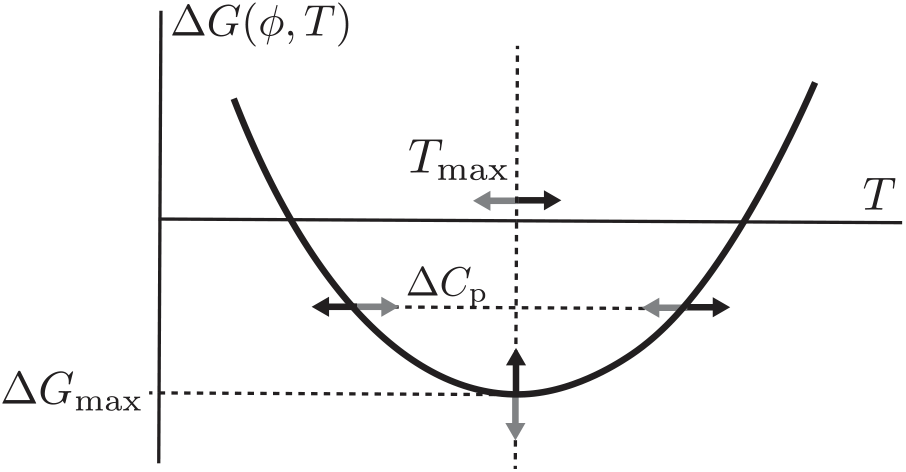
Protein stability curve Δ*G*(***ϕ***, *T*) of a two-state protein as a function of temperature T, parameterized by three thermodynamic parameters ***ϕ*** = (Δ*G*_max_, Δ*C*_p_, *T*_max_), as given in eq. (5). Here, Δ*G*_max_ determines the value of maximum stability (the minimum of the curve), *T*_max_ determines the temperature of maximum stability, and Δ*C*_p_ determines the width of the stability curve. The more negative Δ*G*(***ϕ***, *T*) is, the more stable the protein at T. Black arrows indicate the effect of mutations that increase the corresponding parameter value, while gray arrows indicate mutations that decrease it. The protein stability curve shifts upwards when Δ*G*_max_ increases (decreasing maximum stability), shifts to the right when *T*_max_ increases, and becomes wider when Δ*C*_p_ increases.

We note that the use of the temperature of maximum stability *T*_max_ as the reference temperature is well-established in the literature (Baldwin, 1986; Robertson and Murphy, 1997; Rees and Robertson, 2001; Pucci and Rooman, 2014; Ghosh and Dill, 2009). In contrast to the more commonly used melting temperature, *T*_m_, our choice of using *T*_max_ ensures that the three thermodynamic parameters each have a distinct effect on the protein stability curve (Figure 2). Moreover, similar approximations to ours (eq. 5) have been applied previously as well (Hawley, 1971; Yeritsyan and Badasyan, 2024). While temperature is the primary focus of this work, we acknowledge that other factors, such as pH and salt concentration, can also influence enthalpic interactions and entropic constraints, further modulating protein stability. These additional factors are not considered here.

### Linear model of protein stability

For some proteins temperature-independent enthalpy Δ*H*(***ϕ***) := Δ*H*(***ϕ***, *T*) and entropy Δ*S*(***ϕ***) := Δ*S*(***ϕ***, *T*) fit the data quite well, at least close to their melting temperatures (Barrick, 2018), and we will use this simpler linear model as our reference model. We employ an equivalent parameterization to eq. (4), expressed as,

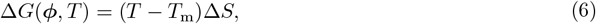

where *T*_m_ = Δ*H/*Δ*S* is the protein’s melting temperature. The two thermodynamic parameters that fully determine protein stability are ***ϕ*** = (Δ*S, T*_m_), and we analyze this model in Sections 3.1.3 and 3.2.5.

#### 2.1.1 Optimal protein stability

A central assumption of this study is that the performance of a phenotype, and hence individual fitness, depends on the stability of its associated protein (eqs. 1-2, see e.g., Fields, 2001; DePristo et al., 2005; Goldstein, 2013; Manhart and Morozov, 2015; Agozzino and Dill, 2018). Proteins that are either overly stable or insufficiently stable become too rigid or too flexible, preventing them from executing their functions efficiently, if at all (Fields, 2001; DePristo et al., 2005; Somero et al., 2017). Consequently, we assume that the performance of the phenotype is maximized at an optimal stability, defined by the probability of the protein being in its folded state (or, alternatively, the proportion of time it remains folded from synthesis to degradation). This optimal probability represents a balance between the protein’s conformational flexibility and rigidity, also referred to as the protein’s optimal fluidity (Somero et al., 2017). The optimal stability can be defined in terms of an optimal equilibrium constant *K*^opt^ = F^opt^*/*U^opt^, where F^opt^ and U^opt^ are concentrations of folded and unfolded proteins at a kinetic equilibrium, respectively, with a normalization F^opt^ + U^opt^ = 1 (Figure 1). They can be alternatively interpreted as the fraction of proteins in their folded and unfolded states, or as the probability of each individual protein to be in its folded and unfolded state. The optimal stability of the protein at temperature *T* can be expressed as

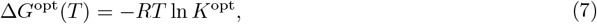

where *R* is the universal gas constant (Figure 1, gray dashed line). Equation (7) is standard in expressing energy in terms of concentration (for ideal gas mixtures; Barrick, 2018). Note that whenever the temperature *T* is assumed constant during viability selection, it can, without loss of generality, be fixed to any arbitrary value because the second parameter, *K*^opt^, can always be scaled without affecting the results.

## 3 Adaptive dynamics of protein stability

In this section, we present results on the evolutionary dynamics of protein stability (all derivations can be found in Appendices A-B and the Supplementary material S1-S2). These dynamics are obtained by performing an invasion analysis of mutations affecting the thermodynamic parameters ***ϕ*** = (Δ*G*_max_, Δ*C*_p_, *T*_max_) (Metz et al., 1992; Geritz et al., 1998; Rousset, 2004). Similar reasoning applies for the linear protein-stability model with ***ϕ*** = (Δ*S, T*_m_) (eq. 6). In this analysis, mutations are assumed to arise rarely, so any prior mutation has either been fixed in the population or has formed a polymorphism with other alleles at a polymorphic equilibrium. As we are interested in the formation of polymorphisms, we assume that initially the population is monomorphic, with all individuals carrying the same resident (wild-type) allele *x*. This allele’s genetic sequence determines the thermodynamic parameters denoted by ***ϕ***_*x*_. We assume throughout that each mutation has only a small effect on these parameters. Thus, for a mutant allele *y* derived from *x*, we write ***ϕ***_*y*_ = ***ϕ***_*x*_ + *d****η***, where each element of the vector ***η*** = (*η*_G_, *η*_T_, *η*_C_) represents the effect of the mutation on parameters Δ*G*_max_, *T*_max_, and Δ*C*_p_, respectively, and *δ* is a small scalar scaling the magnitude of the mutation. Throughout the paper, we assume that Δ*G*_max_ *<* 0, Δ*C*_p_ *<* 0, and *T*_max_ *>* 0 defines the set of biologically feasible values.

The crux of invasion analysis is to calculate the invasion fitness *λ* of *y* in a wild-type population *x*, which determines whether the mutation can invade the population, and to identify and classify so-called singular evolutionary parameters – defined as parameter values where the selection gradient 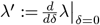 is zero (Geritz et al., 1998). Identifying singular ***ϕ***, which we denoted with ***ϕ***^*^, is central to understanding gradual evolutionary dynamics because the qualitative nature of selection changes only at singularities. Away from ***ϕ***^*^ where the selection gradient is non-zero, selection is always directional and favours mutations yielding a positive selection gradient—any new successfully invading mutant will then substitute the wild-type and fix in the population (Geritz et al., 1998; Dercole and Rinaldi, 2008; Priklopil and Lehmann, 2020). Depending on the sign of selection gradient, mutation-invasion-fixation process thus either moves ***ϕ*** closer to the singular ***ϕ***^*^, in which case we call ***ϕ***^*^ a convergent stable singularity, or moves ***ϕ*** further away from ***ϕ***^*^, in which case we call ***ϕ***^*^ a repellor (REP). When ***ϕ*** is sufficiently close to ***ϕ***^*^, selection can generically only be either stabilizing or disruptive. When selection is stabilizing at a convergent stable ***ϕ***^*^, it is called a convergent stable ESS, also known as a continuously stable strategy (CSS). A CSS can be considered evolutionarily optimal or an evolutionary endpoint, because after ***ϕ*** has converged to the vicinity of ***ϕ***^*^, any new mutation that invades the population will be lost in long term. In contrast, a convergent stable singularity where selection is disruptive is called an evolutionary branching point (EBP), which favors polymorphisms and can thus facilitate local adaptation. The main focus of our analysis is to delineate the conditions under which singular ***ϕ***^*^ are CSSs *versus* EBPs.

### 3.1 Thermally homogeneous environments

Here, we consider a population living in a single habitat with thermal fluctuations around some arbitrary mid-temperature 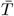.Because we suppose that all individuals in the population experience the same thermal conditions, we refer to this population as the homogeneous population. The population genetics of this model (eqs. 1-3) simplifies to a recursion 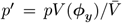,where *p* is the frequency of *y* and 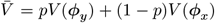 is the average viability in the population. Invasion fitness of *y* in a wild-type population *x* is then *λ* = *V* (***ϕ***_*y*_)*/V* (***ϕ***_*x*_), and the selection gradient becomes 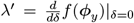 (Appendix B). Below, we will first present results for homogeneous populations that experience small thermal fluctuations around the mid-temperature 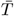,and then in Section 3.1.2 we explore the effect of large thermal fluctuations about 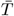.

#### 3.1.1 Small thermal fluctuations

In the model with small thermal fluctuations (*ϵ* = 0) we use 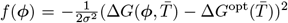 in eqs. (1)-(3), and the singularity 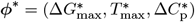 can be solved from

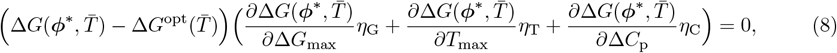

which is obtained by setting 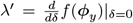 to 0. To calculate ***ϕ***^*^, we will consider each singular 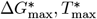 and 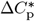 separately by assuming the other two parameters are constant and not affected by mutations (i.e., we set two elements of the vector ***η*** to 0). For now, we note that each singularity can be resolved by setting in eq. (8) either the expression in the first bracket or the second bracket to 0.

##### Adaptive dynamics of maximum stability Δ*G*_max_

Here we assume that mutations affect a single evolutionary parameter, Δ*G*_max_, while the other two thermodynamic parameters, *T*_max_ and Δ*C*_p_, remain constant (i.e., we set *η*_C_ = *η*_T_ = 0 in eq. 8) and fix them at biologically feasible values. Then, recurrent invasions and fixations of new mutations will lead Δ*G*_max_ to reach a CSS maximum stability,

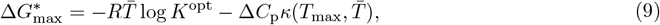

where *κ* is as defined in eq. (5). The CSS maximum stability is given as the difference between the optimal stability of the protein, 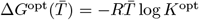 (the first term on the right-hand side), and the temperature-dependent term scaled by heat capacity, 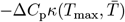 (the second term on the right hand side). We see that selection favors a CSS maximum stability 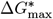 that, for any given parameters *T*_max_, Δ*C*_p_, *K*^opt^, and a mean temperature 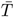,results in a protein stability curve that equals the optimal protein stability, i.e., 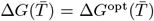. We demonstrate this in Figure 3A by depicting two protein stability curves parameterized by 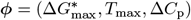,with two different 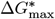 corresponding to two different values of *T*_max_ and Δ*C*_p_.

**Figure 3:**
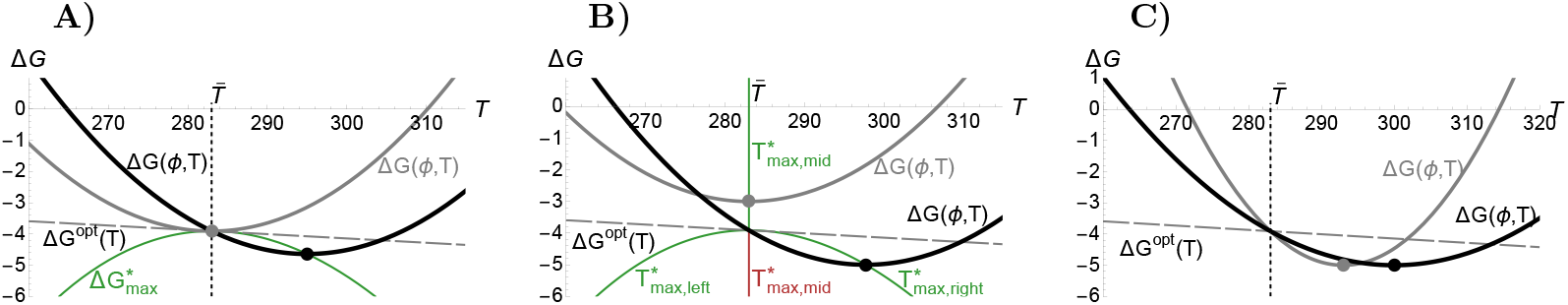
Singular 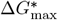 (Panel A), 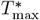 (Panel B), and 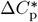 (Panel C) in a homogeneous model with negligible thermal fluctuations (*ϵ* = 0) around a mean temperature 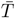,when mutations affect a single parameter at a time. Each panel shows two examples of protein stability curves. The vertical dotted line indicates the mean temperature 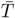 in the habitat, and the descending dashed line represents the optimal stability of the protein Δ*G*^opt^(*T*). In all panels, these two lines intersect at 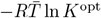 ln *K*^opt^, where 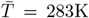 and *K*^opt^ = F^opt^*/*(1 − F^opt^) with F^opt^ = 0.999. **A)** Two protein stability curves for different values of 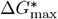 calculated from eq. (9), one where 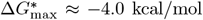 (gray dot), corresponding to (*T*_max_, Δ*C*_p_) = (283K, −3kcal/mol/K), and another where 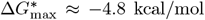 (black dot), corresponding to (*T*_max_, Δ*C*_p_) = (295K, −3 kcal/mol/K). The green dashed curves depict a CSS 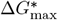 (on the y-axis) for a range of *T*_max_ values (on the x-axis). **B)** Two protein stability curves for different values of 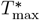 calculated from eq. (10), one where 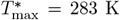 (gray dot), corresponding to (Δ*G*_max_, Δ*C*_p_) = (−3 kcal/mol, −3 kcal/mol/K), and another where 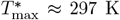 (black dot), corresponding to (Δ*G*_max_, Δ*C*_p_) = (5 kcal/mol, − 3 kcal/mol/K). The color scheme is the same as in Panel A, except that red indicates a repellor. Note that the singular 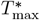 consists of three distinct solutions (eq. 10). **C)** Two protein stability curves for different values of 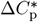 calculated from eq. (11), one where 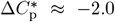 (gray dot), corresponding to (Δ*G*_max_, *T*_max_) = (− 5 kcal/mol, 293K), and another where Δ*C*_p_≈ −5.8 kcal/mol/K (black dot), corresponding to (Δ*G*_max_, *T*_max_) = (−5 kcal/mol, 300K). Unlike 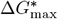 and 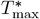 in Panels A and B (green and red dashed lines), we have not depicted 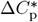 as it has different units than the x- and y-axis.

The effect of model parameters on 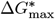 can be obtained from eq. (9). Whenever 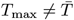,proteins with more negative Δ*C*_p_, or narrower protein stability curves, will favor a greater CSS maximum stability (more negative 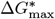,see panel A in Figure 4 for a graphical illustration). This is because, whenever the protein stability curve becomes narrower, the difference between the optimal stability Δ*G*^opt^ and the maximum stability Δ*G*_max_ increases for any fixed *T*_max_, resulting in more negative 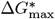.Similarly, a greater deviation between *T*_max_ and the mean temperature 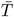 also results in greater CSS maximum stability (Figure 4A). Furthermore, CSS maximum stability increases with increasing *K*^opt^ because the optimal stability Δ*G*^opt^ increases, while the parameter *σ* has no effect because only the optimal value for stability but not the strength of selection impact the value of singularities. The above results imply, for example, that an increasing 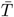 will impose a selective pressure to decrease 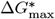 whenever 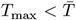,and to increase 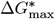 whenever 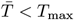 Figure 3A).

**Figure 4:**
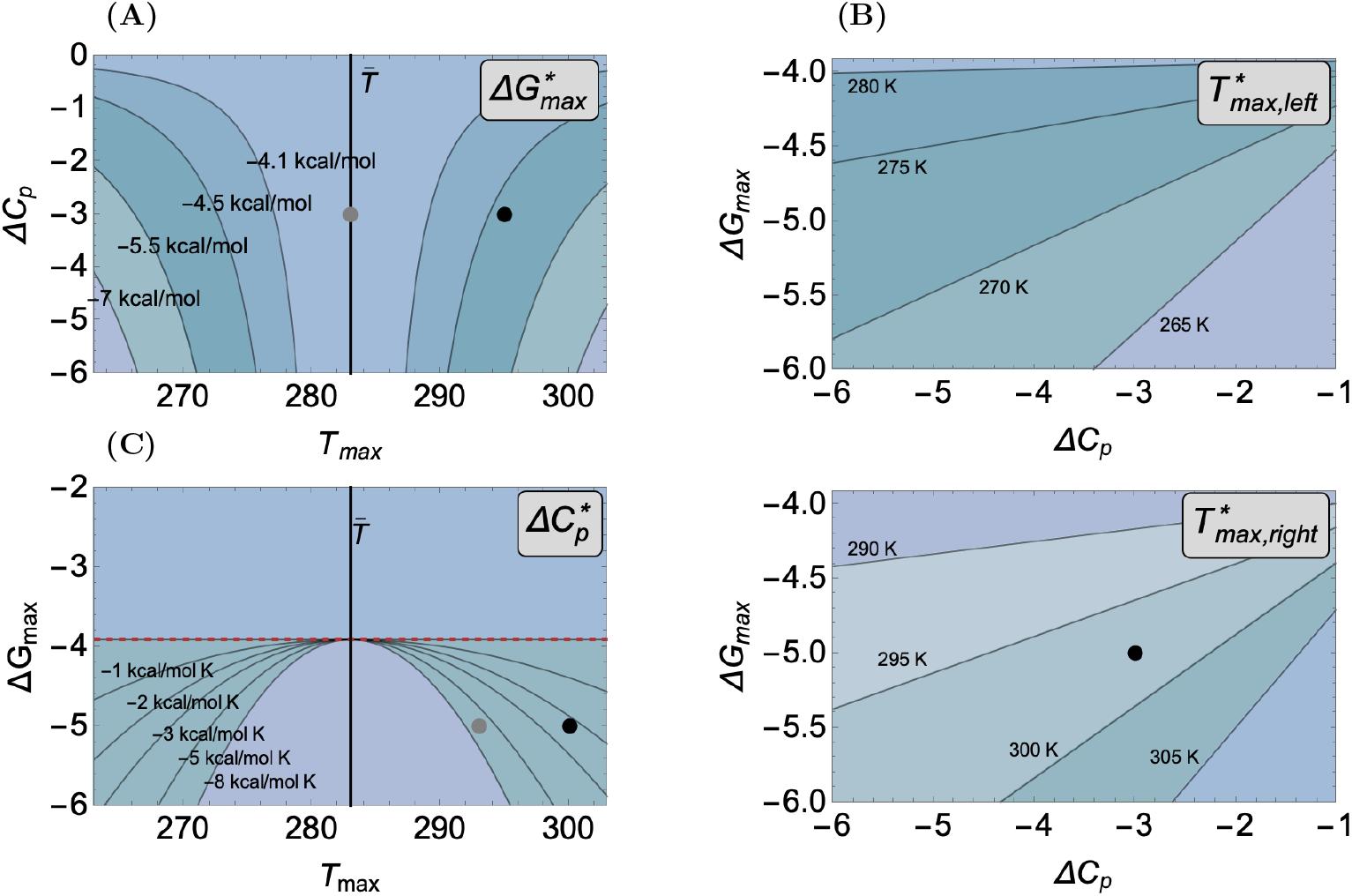
Contours for singular 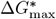 (panel A), 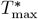 (panels B) and 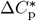 (panel C) as a function of the other two thermodynamical parameters in a homogeneous model without thermal fluctuations (*ϵ* = 0). All depicted singularities are CSSs, and in all panels we use 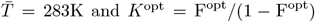 and *K*^opt^ = F^opt^*/*(1 − F^opt^) with F^opt^ = 0.999. **A)** Contours for CSS maximum stability 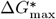 as a function of *T*_max_ and Δ*C*_p_ as given in eq. (10). The contours are nearly symmetric about 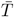.The gray dot at (*T*_max_, Δ*C*_p_) = (283K, − 3kcal/(mol K)) and the black dot at (*T*_max_, Δ*C*_p_) = (295K, − 3kcal/(mol K)) correspond to the black and gray stability curves, respectively, in Figure 3A. **B)** Contours for CSS maximum-stability temperature 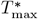 as a function of Δ*C*_p_ and Δ*G*_max_. The top panel shows the left and the bottom panel the right solution as given in eq. (10). In the bottom panel the black dot at (Δ*C*_p_, Δ*G*_max_) = (− 3kcal/(mol K), − 5kcal/(mol K)) corresponds to the black stability curve in Figure 3C. **C)** Contours for CSS heat capacity 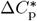 as a function of *T*_max_ and Δ*G*_max_ as given in eq. (11). The contours are symmetric about 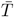.The gray dot at (*T*_max_, Δ*G*_max_) = (293K, −5kcal/(mol K)) and the black dot at (*T*_max_, Δ*G*_max_) = (300K, −5kcal/(mol K)) correspond to the black and gray stability curves, respectively, in Figure 3C. Above the red dashed line where Δ*G*_max_ *>* Δ*G*^opt^, the CSS 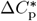 is greater than 0 and hence not biologically feasible.

##### Adaptive dynamics of maximum-stability temperature *T*_max_

Here, we suppose that mutations affect only maximum-stability temperature, *T*_max_, and that the other two thermodynamical parameters, Δ*G*_max_ and Δ*C*_p_, remain constant (i.e., we set *η*_G_ = *η*_C_ = 0 in eq. 8) and fix them at biologically feasible values. We find three biologically feasible singular solutions,

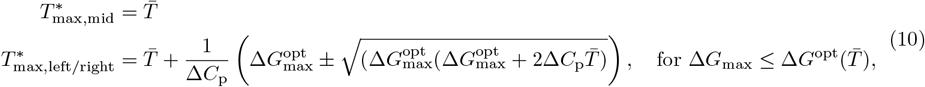

where we use the shorthand notation 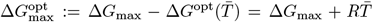 ln *K*^opt^ for the distance between the maximum stability Δ*G*_max_ and the optimal stability 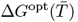.The solutions 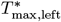 and 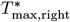 are real-valued and thus biologically feasible only when 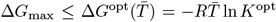. Whenever they are real, they lie on both sides of the mid-solution, with 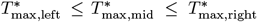,and all three solutions coincide at 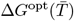 (Figure 3B). The mid-solution 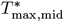 is a CSS whenever 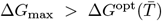,while 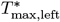 and 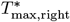 are CSSs, and the mid-solution 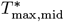 is a repellor (REP) for 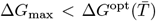 (Figure 3B). This implies that the initial combination of parameter values will determine to which of the three solutions given in eq. (10) the maximum-stability temperature *T*_max_ will converge. When 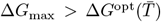,selection favors maximum-stability temperatures that minimize the distance between the protein’s stability 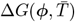 and the optimal stability 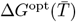.When 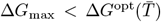,selection favors maximum-stability temperatures for which 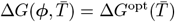. In Figure 3B, we illustrate this by depicting two protein stability curves parameterized by 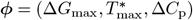 with two different values of Δ*G*_max_ and Δ*C*_p_.

The effect of model parameters on the singularities can be derived from eq. (10) (further details in Supplementary material S1.1 and S2). We observe that only 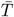 affects the mid-solution 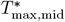.Generally, model parameters have opposite effects on the left 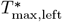 and right solutions 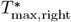.Specifically, 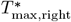 increases (while 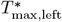 decreases) with increasing Δ*C*_p_ and decreasing Δ*G*_max_. Additionally, 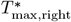 increases (while 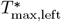 decreases) with decreasing *K*^opt^. However, 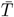 increases, both 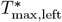 and 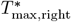 increase. These results imply that an increasing 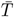 will impose selective pressure to increase 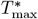 regardless of which branch of solutions the population is adapted to (Figure 3B).

##### Adaptive dynamics of heat capacity Δ*C*_p_

Here we suppose that mutations affect only Δ*C*_p_, and that the other two thermodynamical parameters Δ*G*_max_ and *T*_max_ are constant and fixed at some biologically feasible values (i.e., we set *η*_G_ = *η*_T_ = 0 in eq. 8). We find a unique CSS heat capacity,

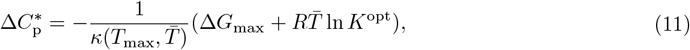

which is negative and hence biologically feasible only when 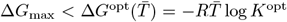.This implies that for 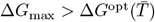 evolution favours ever-increasing values of Δ*C*_p_ until it reaches the boundary of feasible values, where Δ*C*_p_ = 0. The value of the CSS heat capacity 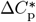 is such that for any given parameters *T*_max_, Δ*G*_max_, *K*^opt^ and 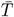,the resulting protein stability is equal to the optimum protein stability at the mean temperature of the habitat, i.e., 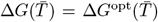.In Figure 3C, we depict two protein stability curves parametrized by 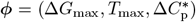 with two different values of (Δ*G*_max_, *T*_max_).

The effect of model parameters on 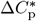 can be obtained directly from eq. (11). The CSS 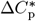 decreases with decreasing Δ*G*_max_ and whenever *T*_max_ gets closer to 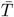 (panel C in Figure 4). Importantly, note that when *T*_max_ approaches 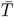, every value of Δ*C*_p_ becomes singular, which we call a degenerate singularity. Moreover 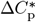 decreases with decreasing *K*^opt^, and whenever 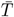 gets closer to *T*_max_, while *σ* has no effect. These results imply that an increasing 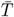 will impose selective pressure to decrease 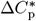 whenever 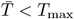 and to increase 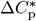 whenever 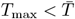 (Figure 3C).

##### Adaptive dynamics of the vector *ϕ* = (Δ*G*_max_, *T*_max_, Δ*C*_p_)

In the preceding paragraphs, we derived analytical expressions for the singular solutions 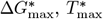 and 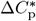 in a population experiencing negligible thermal fluctuations around a mean temperature 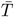,and analyzed their evolutionary properties when mutations affect one parameter at a time. Here, we briefly consider what happens if mutations sequentially affect each parameter as this reveals the possible adaptive dynamics of the entire stability profile ***ϕ***. From eq. (8), we observe that once Δ*G*_max_ has converged to the singular value 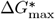 (given in eq. 9), the other two parameters, *T*_max_ and Δ*C*_p_, also satisfy the singularity condition (eq. 8) and hence become singular. This occurs because a singular 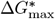 is uniquely determined by setting the first bracket in eq. (8) to zero (since ∂Δ*G*(***ϕ***)*/*∂Δ*G*_max_ = 1), which is the singularity condition for *T*_max_ and Δ*C*_p_ as well. Similarly, when Δ*C*_p_ converges to 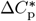 (as given in eq. 9), which is negative and thus biologically feasible only if 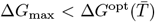,the other two parameters, *T*_max_ and Δ*G*_max_, also become singular. Finally, when *T*_max_ converges to 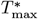 (as given in eq. 9), Δ*G*_max_ and Δ*C*_p_ also become singular. However, if 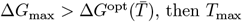,then *T*_max_ converges to 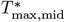,in which case Δ*C*_p_ becomes a selectively neutral singularity. Therefore, caution is required when analyzing the convergence stability of the vector-valued trait ***ϕ***, as the mutation process can influence which type of singularity the population converges to (Leimar, 2005, 2009). From this discussion, we conclude that there are generally three qualitatively different outcomes. If 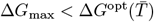,then regardless of the mutation process, the vector ***ϕ*** will converge to the CSS ***ϕ***^*^, which is always biologically feasible. In this scenario, *T*_max_ evolves to either 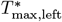 or 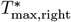,and 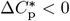.The evolutionary endpoint is then given by eqs. (9)–(11). However, if 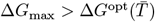,then depending on the mutation process, ***ϕ***^*^ can either be biologically feasible, as previously described, or *T*_max_ converges to 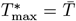, and Δ*C*_p_ either converges to the boundary of its feasible values or remains neutral and unaffected by selection. We will address the analysis of multivariate mutations and the effects of trade-offs in later sections, as trade-offs can lead to polymorphisms. These polymorphisms are relevant to local adaptation, which is the focus of Section 3.2.

#### 3.1.2 Large thermal fluctuations

Here we analyse the effect of non-negligible thermal fluctuations (*ϵ >* 0) around the mid-temperature 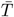 on the evolutionary dynamics of Δ*G*_max_, *T*_max_ and Δ*C*_p_. First, we suppose that mutations affect a single evolutionary parameter Δ*G*_max_, and that the other two thermodynamical parameters, *T*_max_ and Δ*C*_p_, remain constant and fixed at some arbitrary but biologically feasible values. The singular 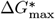 is given as

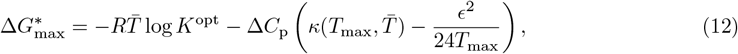

where the term in the brackets is the average annual effect of temperature on protein stability. This term is composed of two terms, the effect of temperature on protein stability at the mean temperature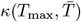,and of a correction term *ϵ*^2^*/*(24*T*_max_) due to the thermal fluctuations acting on *κ*. Similarly to the model with negligible thermal fluctuations, the singularity 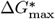 (eq. 12) is always a CSS.

The effect of thermal fluctuations comes through the correction term *ϵ*^2^*/*(24*T*_max_) that was absent in the model with negligible thermal fluctuations (compare with eq. 9). Increasing thermal fluctuations thus increase the CSS maximum stability by decreasing 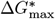,but they only have a minor effect because for biologically realistic parameter values 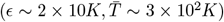 the correction term is small and of the order *O*(10^−1^). This is illustrated graphically in Figure 5A where we show how 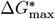 changes as a function of *ϵ* when Δ*C*_p_ = −3kcal/mol and for three different values of *T*_max_. The effect of the mean temperature 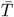 is similar to the model with negligible thermal fluctuations *ϵ* = 0. Greater maximum stability is favoured (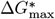decreases) with increasing 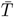 whenever 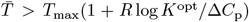,otherwise with decreasing 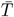.We note that for biologically realistic parameter values this threshold is approximately 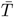,and thus close to the threshold for a model with negligible thermal fluctuations *ϵ* = 0.

**Figure 5:**
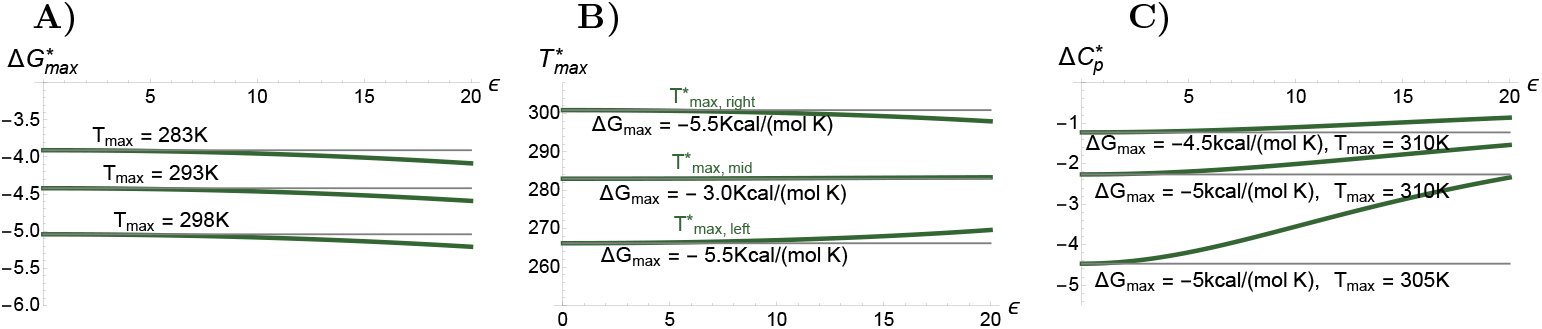
The effect of thermal fluctuations *ϵ* on the singular 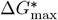 (panel A), 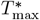 (panel B) and 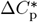 (panel C). The green curves indicate the singularities calculated for different parameter values as a function of *ϵ*, and for comparison, the thin gray straight lines indicate the singularities for the same parameters but in a model where *ϵ* = 0. Note that these solutions coincide as *ϵ* approaches 0. **A)** Singular 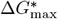 calculated from eq. (12), using 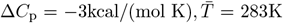 and three different values of *T*_max_. We observe that *ϵ* has only a small effect on 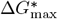.**B)** Singular 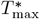 calculated numerically, using 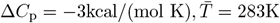 and two different values of Δ*G*_max_. Note that the three curves are for the three different branches of solutions. We observe that *ϵ* has only a small effect on 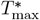.**C)** Singular 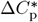 calculated from eq. (13), using 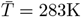 and three different values of (Δ*G*_max_, *T*_max_). We observe that *ϵ* has a relatively large effect on 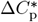.In particular, we observe that either by increasing maximum stability (decreasing Δ*G*_max_) or decreasing *T*_max_ the effect of *ϵ* is larger.

The expressions for the singular 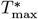 are not presented here because they are lengthy and difficult to interpret. However, our numerical analysis suggests that singular solutions are qualitatively and quantitatively similar to the case with negligible thermal fluctuations (Supplementary material S1.1.1 and S2). That is, above a certain critical threshold of Δ*G*_max_, the only CSS is approximately 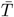,and below this threshold this mid-temperature is a repellor and two branches on both sides of this repellor are the CSSs. In Figure 5B we plot the effect of *ϵ* on the three CSS solutions, suggesting that thermal fluctuations have a small role in determining 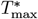.

The singular 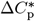 is always a CSS and can be expressed as

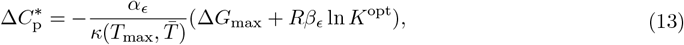

where the coefficients *α*_*ϵ*_, *β*_*ϵ*_ are due to the thermal fluctuations and depend on 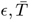 and *T*_max_ (for analytical expressions, see Supplementary material S1.1.2). The coefficients are positive for realistic parameter values, yet, their dependence on *ϵ* is complex and we could only analyse the effect of *ϵ* in the special case where *T*_max_ is close to 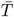.In this special case Δ*C*_p_ increases with increasing *ϵ*, and our numerical analysis suggests that this is also true for the remaining branches of solutions of *T*_max_. Importantly, and in contrast to 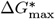 and 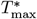,the effect of *ϵ* on 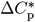 is significant, and even small changes in *ϵ* can have profound effect on 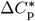 (Figure 5C).

In summary, we find that the model with non-negligible thermal fluctuations (*ϵ >* 0) can be approximated well with the model with negligible thermal fluctuations (*ϵ* = 0), at least with regards to the parameters Δ*G*_max_ and *T*_max_. However, heat capacity Δ*C*_p_ is sensitive to *ϵ*, and the two models depart the most with decreasing values of Δ*G*_max_ (further away it is from Δ*G*^opt^) and closer is *T*_max_ to 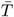.Overall, this suggests that changes in the magnitude of thermal fluctuations has little impact on Δ*G*_max_ and *T*_max_ but yields evolutionarily sub-optimal heat capacity and strong selection acting on Δ*C*_p_.

#### 3.1.3 Linear model of protein stability

Here, we compare the above results with a model in which protein stability varies linearly with temperature (eq. 6). Our analysis focuses on mutations that specifically affect *T*_m_ through changes in Δ*H*. This approach is based on the key objective of this work, which is to understand thermal adaptation to new environments; hence, mutations affecting *T*_m_ are most relevant because they shift protein stability profiles along the temperature axis.

We take into account both the mean 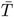 and thermal fluctuations *ϵ* and find that there is a unique CSS expressed as

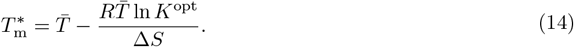

The CSS 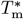 is independent of thermal fluctuations because both the optimal stability and protein stability vary linearly with temperature. Consequently, fluctuations above and below the mean temperature cancel out, making the CSS dependent only on the mean temperature 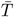.

### 3.2 Thermally heterogeneous environments

In this section we consider a population subdivided into two habitats, L and H, with mid-temperatures 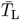 and 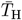,respectively. We will separately examine the evolutionary dynamics of each evolutionary thermodynamic parameter Δ*G*_max_, *T*_max_ and Δ*C*_p_ (Sections 3.2.1-3.2.3). For each parameter, we first focus on small thermal fluctuations (*ϵ* = 0) and calculate their singular values and their evolutionary properties, that is, whether they are CSSs or EBPs. Second, for each parameter, we characterise their adaptive dynamics in two typical biological scenarios. In the first scenario, the population is initially assumed to be adapted to the colder habitat L. A warmer habitat H then becomes available, creating a heterogeneous thermal environment consisting of the two habitats connected by migration. In the second scenario the population is assumed to be initially adapted to the warmer habitat H after which the colder habitat L opens up. For both biological scenarios, we discuss the conditions that favour either global or local adaptation of protein stability. Third, we conclude each section with a discussion on the effect of thermal fluctuations. In Section 3.2.4, we consider the joint adaptive evolution of multiple thermodynamic parameters. Given that their joint evolution is bound to be constrained by biophysical trade-offs between these parameters, we analyze the joint evolution of each pair of parameters and characterize the types of trade-off functions that favor local versus global adaptation of protein stability. Finally, in Section 3.2.5, we compare our results to the simpler linear model (eq. 6).

#### 3.2.1 Adaptive dynamics of maximum stability

Suppose the thermal fluctuations are negligible (*ϵ*_L_ = *ϵ*_H_ = 0) and that mutations affect only the maximum stability Δ*G*_max_. The other two parameters *T*_max_ and Δ*C*_p_ remain constant (i.e., we set *η*_G_ = *η*_T_ = 0) and fix them at some feasible values. We find a unique convergent stable singular maximum stability,

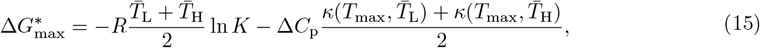

which is always negative and hence biologically feasible. Note that 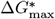 (eq. 15) has a similar form as for homogeneous populations (eq. 9), except of the term 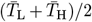,which is the mean temperature across habitats, and 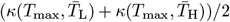 which is the mean effect of temperature on stability with the average calculated across habitats.

The convergent stable 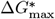 (eq. 15) is a CSS whenever

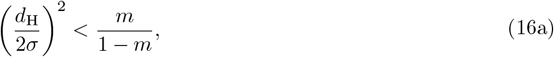

where

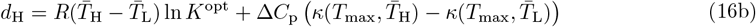

is the sum of the distance between the optimal stability and protein stability in each habitat, that is, 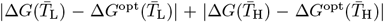.If the CSS condition (eq. 16) is not satisfied, the singularity is an evolutionary branching point EBP. EBP is favoured (CSS condition is disfavoured), whenever the strength of selection *d*_H_*/σ* increases. This will occur, in our model, whenever *σ* decreases or whenever the distance *d*_H_ increases. In both situations any trait that is favoured in one habitat will be strongly selected against in the other habitat leading to disruptive selection and evolutionary branching (local adaptation via gradual evolution). Branching is thus favoured for smaller values of *σ*, and for larger values of *d*_H_ which is the case for larger *K*^opt^ and for smaller Δ*C*_p_. The effect of mean temperatures 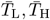 on *d*_H_ is a bit more complex. The distance *d*_H_ increases with increasing 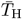 whenever 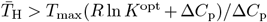,otherwise with decreasing 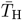.In contrast, the distance *d*_H_ increases with decreasing 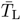 whenever 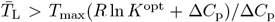,otherwise with increasing 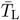.This implies that an increase in the difference between the mean temperatures 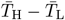 is not sufficient to conclude whether stabilising or disruptive selection is favoured. Note, that for biologically reasonable values *T*_max_(*R* ln *K*^opt^ + Δ*C*_p_)*/*Δ*C*_p_ ≈ *T*_max_, and so EBP is favoured upon an increase (decrease) in 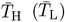 whenever 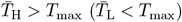.Finally, a decrease in symmetric migration *m* favours EBP and is consistent with the notion that limited gene flow favours local adaptation (Lenormand, 2002; Kawecki and Ebert, 2004, for asymmetric migration, see Priklopil, 2025).

##### Population initially adapted to habitat L

In this biological scenario, we assume that the population initially inhabits and is adapted to habitat L with a mean temperature 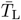,and upon forming a heterogeneous thermal environment with habitat H where 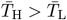,we study the adaptive evolution of Δ*G*_max_. The evolutionary parameters ***ϕ*** are thus initially given as in eqs. (9)-(11). However, as there are infinitely many possible CSS solutions for a homogeneous population adapted to temperature 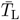,we focus on the case where 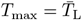 for simplicity. Specifically, we assume that in the initial homogeneous population 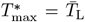,but we drop the asterisk as this singularity is no longer singular in the newly formed heterogeneous environment. Since for this case Δ*C*_p_ is neutral and every value is singular, we use Δ*C*_p_ as a bifurcation parameter.

In Figure 6A-1 we summarize our results on local and global adaptation by showing the evolutionary properties of the convergent stable singularity (eq. 15) as a function of 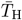 and Δ*C*_p_. We have assumed, without loss of generality, that 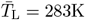 (approximately 10 degrees Celsius). We find that increasing 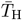 and decreasing Δ*C*_p_ favors disruptive selection and thus the condition for EBP (red area in Figure 6A-1). Moreover, we confirm that increasing *m* indeed favors CSS (blue area), as shown by using two different values of *m*. The contour between the red and blue areas represents the result for *m* = 0.01, while the dashed line shows where the contour would be for *m* = 0.1.

**Figure 6:**
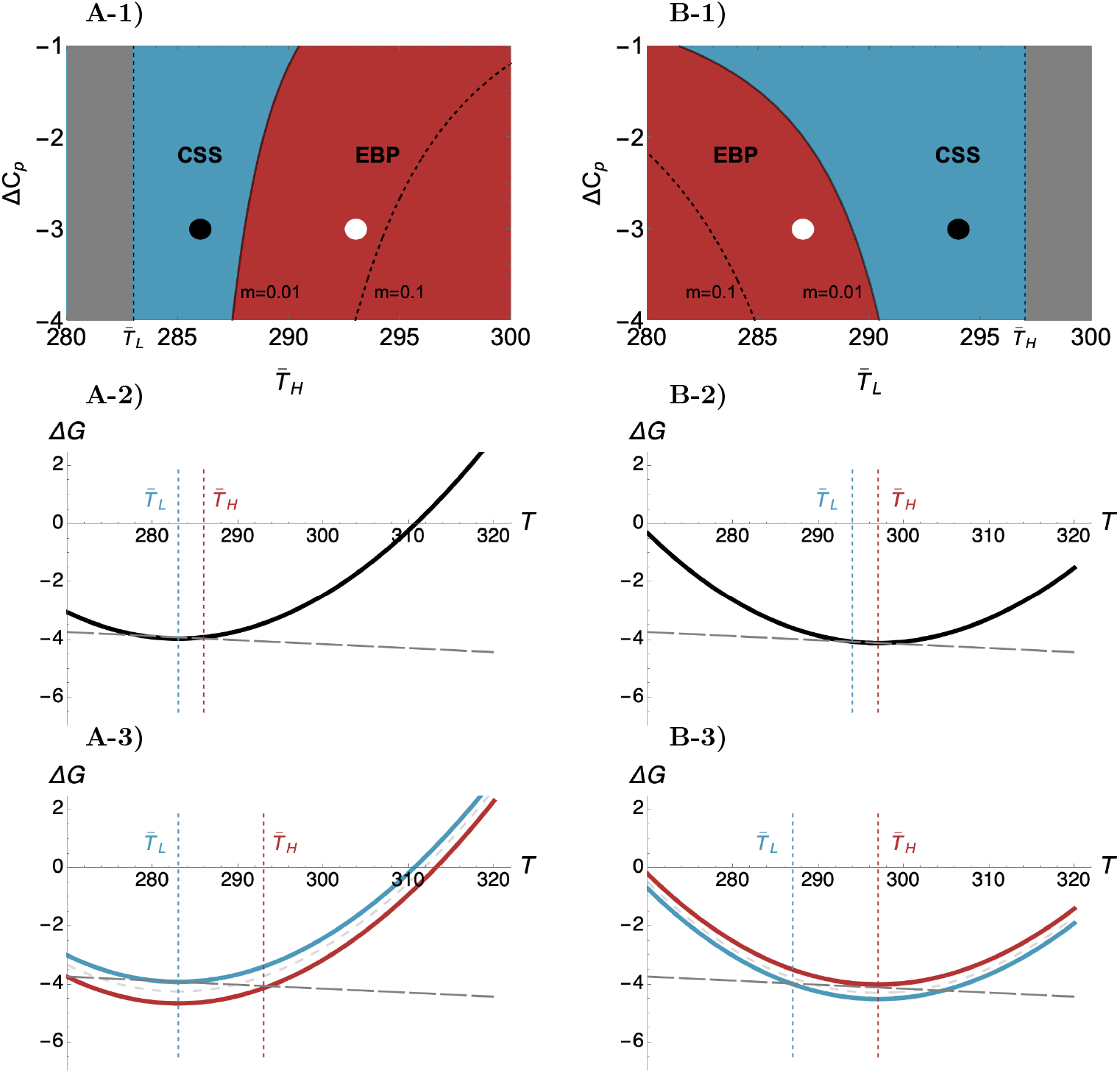
The top panels depict the evolutionary properties of 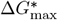 as a function of Δ*C*_p_ and 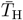 or 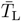,while the middle and bottom panels illustrate instances of globally and locally adapted protein stability curves in the two biological scenarios discussed in the main text (Section 3.2.1). In panels A, we assume 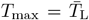,and in panels B, we assume 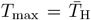.In the top panels, blue indicates that 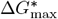 is a CSS, and red indicates that it is an EBP according to the condition in eq. (16). The migration rate is *m* = 0.01, and the dashed line within the red region shows where CSS and EBP would delineate for *m* = 0.1. The middle and bottom panels depict protein stability curves corresponding to the black and white dots in the top panels, respectively. **A-1)** We observe that EBP is favoured with increasing 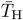 and decreasing Δ*C*_p_. The gray region where 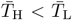 is not biologically feasible. **B-1)** We observe that EBP is favored with decreasing 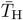 and Δ*C*_p_. This case favors CSS more compared to panel A-1. The gray region where 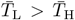 is not biologically feasible. **A-2)** Monomorphic CSS protein stability that is globally adapted across the two thermal environments. The parameter values are 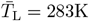 and 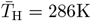,corresponding to the black dot in the top panel A-1. **B-2)** Monomorphic CSS protein stability that is globally adapted across the two thermal environments. The parameter values are 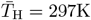 and 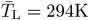,corresponding to the black dot in the top panel B-1. **A-3)** Polymorphic CSS protein stability curves that are locally adapted to the habitat L (blue curve), and habitat H (red curve). The parameter values are 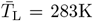 and 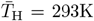,corresponding to the white dot in the panel A-1. The gray dashed curve indicates the singularity that is an EBP and hence only a transient state. **B-3)** Polymorphic CSS protein stability curves that are locally adapted to the habitat L (blue curve) and habitat H (red curve). The parameter values are 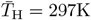 and 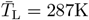,corresponding to the white dot in the panel B-1.

The next question is what kind of protein stability curves are expected to arise as a result of global and local adaptation. We address this question by analysing two instances of adaptation: one where the singularity 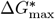 is a CSS (black dot in Figure 6A-1, with 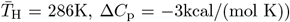 and another where the singularity is an EBP, thus only a transient state leading to polymorphism (white dot in Figure 6A-2, with 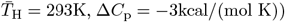,When the singularity is a CSS, the maximum stability Δ*G*_max_ will converge through mutations and fixations to a neighborhood of the singular 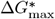 as given in eq. (16), and the population remains monomorphic with respect to this parameter. While new mutations can invade and form a polymorphism with this singular allele, this is only short-term as eventually an allele whose value is even closer to 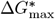 will invade and fix into the population. The resulting protein stability curve is depicted in Figure 6A-2. We observe that the globally adapted 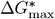 results in a curve whose stability in L is slightly larger than the optimal stability in this habitat 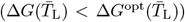 and whose stability in H is slightly smaller than the optimal stability in this habitat 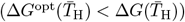.

In contrast, when the singularity 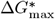 is an EBP, the maximum stability Δ*G*_max_ will also converge through mutations and fixations to a neighborhood of the singular 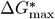 as given in eq. (15), but then experiences disruptive selection, causing the population to become polymorphic with respect to this parameter. Subsequent mutations and invasions will tune the polymorphism to a vicinity of a polymorphic CSS with two locally adapted alleles. Similarly to the previous case, new mutations can invade and form a polymorphism of three alleles, but this is only short-term as eventually a pair of alleles whose values are even closer to the polymorphic CSS will invade and fix into the population. The resulting protein stability curves are depicted in Figure 6A-3. We observe that the allele adapted to L results in a curve with maximum stability close to the optimal stability in L, i.e., 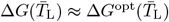,and the allele adapted to H results in a curve that intersects close to the optimal stability in habitat H, i.e., 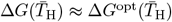.We conclude that in this biological scenario, local adaptation of protein stability is achieved by mutations that increase the maximum stability of the protein stability curve initially adapted to habitat L, i.e., by decreasing Δ*G*_max_.

##### Population initially adapted to habitat H

In this biological scenario, we suppose that the population is initially inhabited and adapted to habitat H with mean temperature 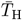.Then, upon forming a heterogeneous thermal environment with habitat L where 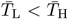,we study the adaptive evolution of Δ*G*_max_. Analogously to the above biological scenario, we suppose that 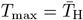 and use Δ*C*_p_ as a bifurcation parameter. In Figure 6B-1, assuming 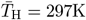,we show the evolutionary properties of the convergent stable singularity (eq. 15) as a function of 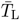 and Δ*C*_p_. We find that decreasing 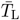 and Δ*C*_p_ favors disruptive selection and thus leads to the condition for EBP. Similarly to the previous scenario, EBP is favored with decreasing *m*.

To investigate the global and local adaptation of protein stability curves, we again present two instances of adaptation. One where the singularity 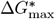 is a CSS (black dot in Figure 6A-1, with 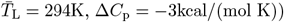 and another where the singularity is an EBP, thus only a transient state leading to polymorphism (white dot in Figure 6A-2, with 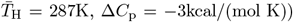.We observe a similar pattern of local and global adaptation as in the previous scenario. Specifically, local adaptation of protein stability is achieved by mutations that increase the maximum stability, i.e., by decreasing Δ*G*_max_. This is interesting as the population is adapting to a habitat with a lower mean temperature and hence lower optimal stability. Yet, this is achieved by mutations that increase the maximum stability of the protein stability curve initially adapted to habitat H.

Comparing the two biological scenarios, we observe that the condition for EBP is more stringent for this case compared to if the population was initially adapted to L. This is because the same difference in temperature in the two habitats imposes different selection pressures in the two cases. This is a consequence of the fact that the optimal protein stability is smaller in L than in H, and that protein stability curves open upwards. As a result, the difference between protein stability curves adapted to L and H is smaller, leading to stronger stabilizing selection if the population was initially adapted to H where 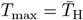.This is evident when comparing Figures 6A-3 and B-3, as the difference between the maximum stabilities adapted to the two different habitats is larger in A-3 than in B-3. Adaptation in the second biological scenario is thus expected to require fewer mutations and is thus potentially faster.

##### Large thermal fluctuations

If thermal fluctuations *ϵ*_L_ and *ϵ*_H_ around the mid-temperatures 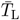 and 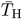,respectively, are non-negligible, the convergent stable singularity takes the form

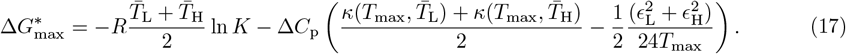

Because Δ*C*_p_ is negative, increasing thermal fluctuations thus lead to an increase in singular protein stability (decrease in 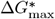). This occurs because the protein stability curve deviates most from optimal stability at extreme temperatures. As a result, to minimize fitness costs at the most extreme temperatures in each habitat, the protein stability curve shifts downward. The convergent stable 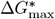 (eq. 17) is a CSS whenever eq. (16a) is coupled with

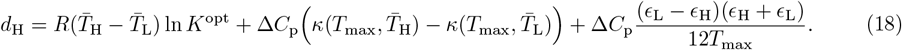

We observe that thermal fluctuations have only a minor effect on evolutionary stability because the denominator in the final term is several orders of magnitude larger than the nominator. Moreover, the effect of fluctuations cancels out entirely if the fluctuations in both habitats are identical (*ϵ*_L_ = *ϵ*_H_). This is due to the fact that when 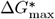 is singular, the average fitness costs in each habitat is the same regardless of the magnitude of fluctuations. Since Δ*G*_max_ can only shift the protein stability curve up or down, there is no fitness benefit in specializing for one habitat over the other and hence does not affect evolutionary stability.

#### 3.2.2 Adaptive dynamics of the maximum-stability temperature

Here, we assume that mutations affect the maximum-stability temperature *T*_max_, while the other two parameters, Δ*G*_max_ and Δ*C*_p_, remain constant (i.e., *η*_G_ = *η*_C_ = 0) and fixed at some feasible values. Unfortunately, we could not find concise analytical solutions for this biological scenario. Numerical explorations however indicate that there are up to 4 solutions with complex patterns of evolutionary stability, rendering this case out of scope of this paper. We thus focus on the numerical analysis of the two special biological scenarios.

##### Population initially adapted to habitat L

In this biological scenario we suppose that the population initially inhabited habitat L, and that the maximum stability was adapted to this habitat with 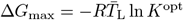 while Δ*C*_p_ is neutral and used as a bifurcation parameter. We could not find a concise analytical expression for singular values, but in Supplementary material S1.2.1 we show that there exists a single convergent stable singularity, and that this singularity lies in the interval 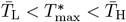.Moreover, by using the method of implicit differentiation on the condition for singularity (see e.g., Otto and Day, 2011, Section 12.3), we find that this singularity increases with increasing 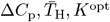,and that *σ* and *m* have no effect. In Figure 7A, we assume 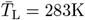 and show the evolutionary properties of the convergent stable singularity as a function of 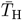 and Δ*C*_p_. We observe a similar result than for the evolution of Δ*G*_max_ (Section 3.2.1), in that increasing 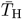 and decreasing Δ*C*_p_ favours disruptive selection and hence the condition for EBP (red area in Figure 7A). Moreover, also increasing *m* favours CSS (blue colour), which we depict by using two different values of *m*, the contour between red and blue colour gives the result for *m* = 0.01 and the dashed line shows where the contour between red and blue would be for *m* = 0.1.

**Figure 7:**
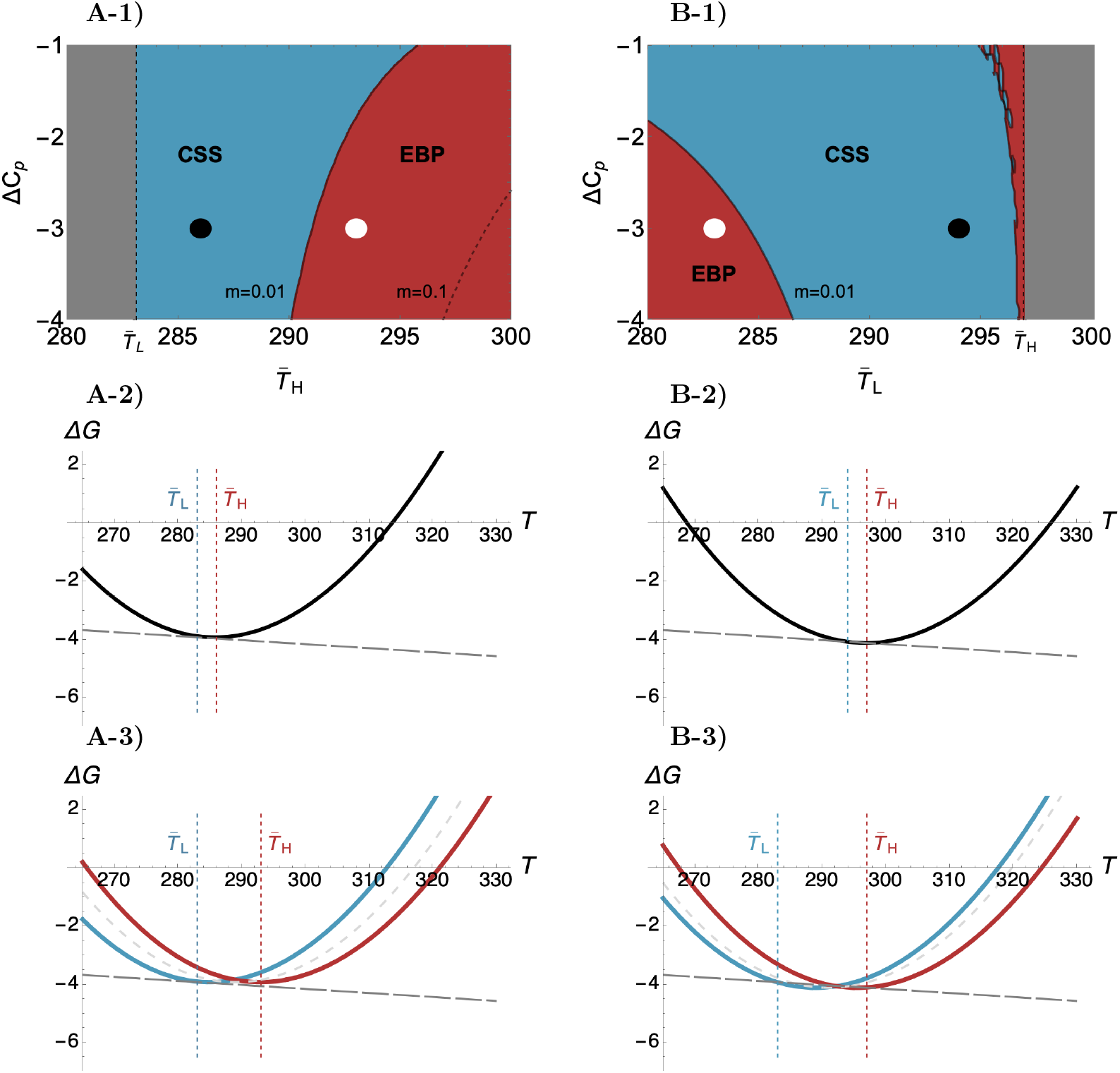
The top panels depict the evolutionary properties of 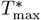 as a function of Δ*C*_p_ and 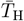 or 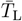,while the middle and bottom panels illustrate instances of globally and locally adapted protein stability curves in the two biological scenarios discussed in the main text (Section 3.2.2). The structure and the color scheme of the panels are identical to Figure 6. **A-1)** We observe that EBP is favoured with increasing 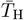 and decreasing Δ*C*_p_. The gray region where 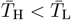 is not biologically feasible. **B-1)** We observe that EBP is favoured with decreasing 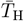 and Δ*C*_p_. Note that this case favours CSS as compared to the panel A-1. The gray region where 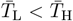 is not biologically feasible. **A-2)** Monomorphic CSS protein stability that is globally adapted across the two thermal environments. The parameter values are 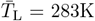 and 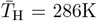,corresponding to the black dot in the top panel A-1. **B-2)** Monomorphic CSS protein stability that is globally adapted across the two thermal environments. The parameter values are 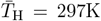 and 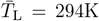,corresponding to the black dot in the top panel B-1. **A-3)** Polymorphic CSS protein stability curves that are locally adapted to the habitat L (blue curve), and habitat H (red curve). The parameter values are 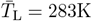 and 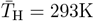,corresponding to the white dot in the top panel A-1. **B-3)** Polymorphic CSS protein stability curves that are locally adapted to the habitat L (blue curve), and habitat H (red curve). The parameter values are 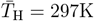 and 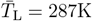,corresponding to the white dot in the top panel B-1.

##### Population initially adapted to habitat H

In this biological scenario we suppose that the population initially inhabited habitat H, and the maximum stability was adapted to this habitat with 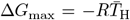 ln *K*^opt^. In contrast to the previous case, however, singular solutions 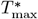 outside of the interval between 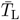 and 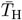 are possible, and, we can not establish their convergence stability. Never-theless, we can investigate how the singular *T*_max_, whatever its value, is affected by parameters. We find that 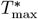 increases with increasing 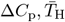 and *K*^opt^, and that *σ* and *m* have no effect. In Figure 7B, we assume 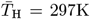 and show the evolutionary properties of the convergent stable singularity as a function of 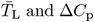 and Δ*C*_p_. We observe a similar result than for the evolution of Δ*G*_max_, in that decreasing 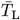 and decreasing Δ*C*_p_ favours disruptive selection and hence the condition for EBP (but note that we encountered minor numerical issues for 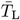 values close to 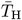;see the rugged region in Figure 7B-1). Moreover, increasing *m* favours CSS (blue colour). We also note that similarly to the case of Δ*G*_max_, disruptive selection is weaker in this case as compared to the case where population was initially adapted to L.

Comparing the two biological scenarios we observe that similarly to Δ*G*_max_ (Section 3.2.1), the condition for EBP is more stringent if the population is adapted to H than to L. This is because the protein stability curve that was pre-adapted to H has a greater maximum stability (smaller Δ*G*_max_) than if it were pre-adapted to L, and hence all else being equal, a smaller change in the parameter *T*_max_ is needed for the stability to adapt to the alternate habitat in the former case. This is evident when comparing the two curves in panels B-3 and A-3 (Figure 7), as in the former case (panel B-3) the two stability curves are closer to each other. Adaptation in the scenario where population is initially adapted to H is thus expected to require fewer mutations and is hence potentially faster.

##### Large thermal fluctuations

We will now briefly explore the role of thermal fluctuations in the two biological scenarios discussed above. For simplicity, we focus on equal thermal fluctuations in each habitat, *ϵ*_L_ = *ϵ*_H_ = *ϵ*. In Figure 8, we present the evolutionary properties of the unique convergent stable singular 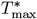.In both biological scenarios (panels A and B), we find that while thermal fluctuations have a minimal effect on the singularity, which remains approximately midway between 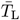 and 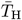 (not shown), they have a major impact on evolutionary stability. That is, increasing thermal fluctuations favours evolutionary branching and thus local adaptation. This occurs because, at extreme temperatures in each habitat, protein stability deviates most from the optimal stability, leading to greater fitness costs as thermal fluctuations increase. These fitness costs can be minimized if one allele specializes in one habitat and another allele specializes in the other, thereby favoring evolutionary branching. Interestingly, the arguments regarding the effects of thermal fluctuations on the singularity and the CSS conditions are exactly opposite to those made for Δ*G*_max_ in Section 3.2.1. This makes sense, as Δ*G*_max_ and *T*_max_ have orthogonal effects on protein stability.

**Figure 8:**
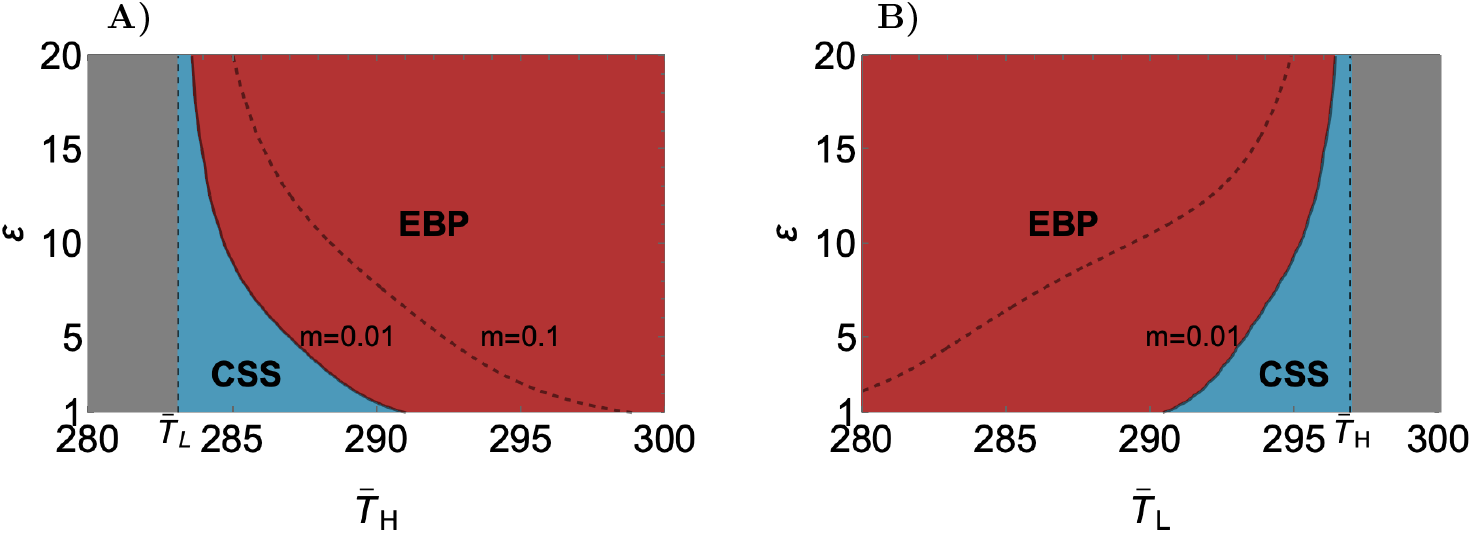
A plot of the evolutionary properties of 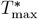 as a function of thermal fluctuations *ϵ* and 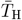 (panel A) 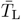 (panel B). The color scheme is identical to Figure 6. In both panels A and B, increasing thermal fluctuations and the across habitat difference in temperature favour evolutionary branching. **A)** In this panel we have assumed that 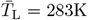 and Δ*C*_p_ = −3kcal/(mol K). **B)** In this panel we have assumed that 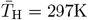 and Δ*C*_p_ = −3kcal/(mol K).

#### 3.2.3 Adaptive dynamics of heat capacity

Here we suppose that mutations only affect heat capacity Δ*C*_p_, while the other two evolutionary parameters Δ*G*_max_ and *T*_max_ remain constant (i.e., we set *η*_G_ = *η*_T_ = 0), and fix them at some feasible values. We find that there exists a unique convergent stable singularity,

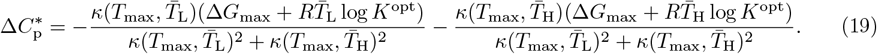

Depending on the combination of parameter values, this solution can be negative or positive and hence can lie outside the set of feasible values (in Figure 9, positive 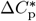 is indicated with gray colour). Unfortunately, the condition for evolutionary stability in this general biological scenario is lengthy, and so we provide numerical analysis. In Figure 9 we depict the parameter combinations (*T*_max_, Δ*G*_max_) for which the singularity is a CSS or EBP. In panel 9A we assume 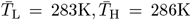,and in panel 9B we assume 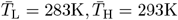.Interestingly, larger difference between the habitat-specific temperatures 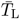 and 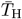 does not appear to favour EBP, as the condition for EBP (red regions) is satisfied for a considerable range of parameter values in both cases. CSS is notably favoured when *T*_max_ is close to either of the mid-temperatures 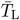 or 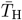 (we confirm this analytically below). This is because for this case, when Δ*G*_max_ is more negative than the optimal stability, one can adjust the width of the curve so that it is well adapted across the two environments. However, if *T*_max_ lies in the interior 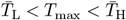 (but not right in the middle, see panel B), then two different widths are favoured hence also favouring EBP.

**Figure 9:**
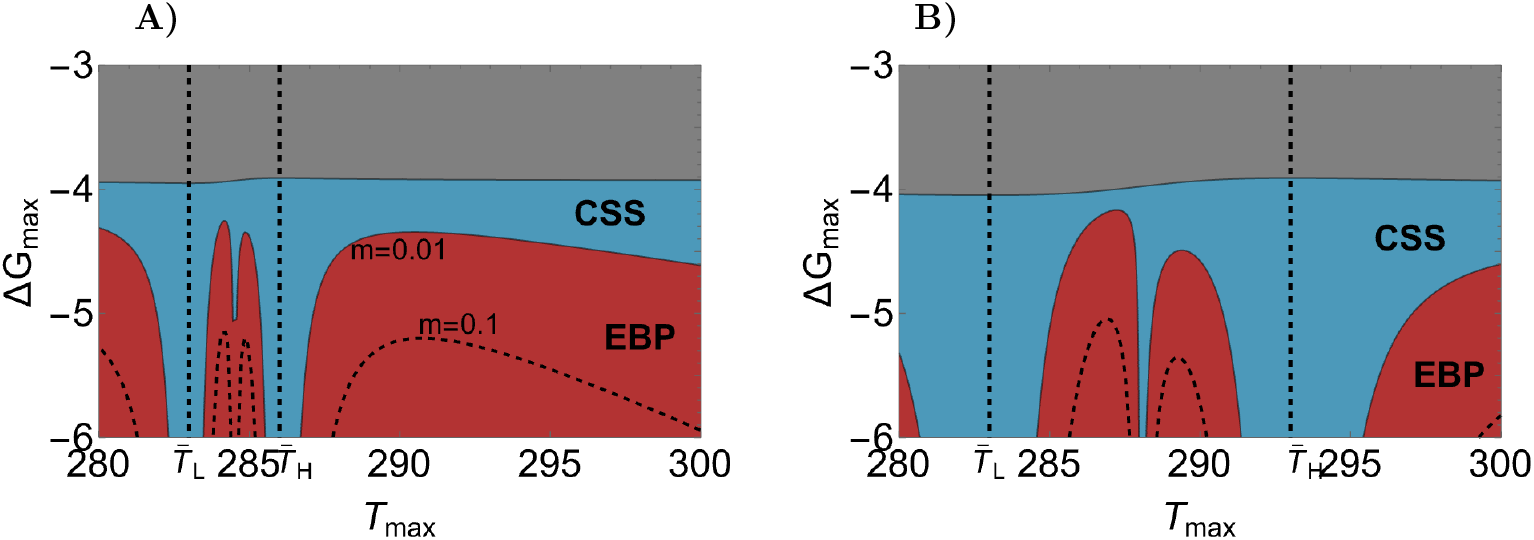
A plot of the evolutionary properties of 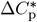 as a function of *T*_max_ and Δ*G*_max_ in a general biological scenario (Section 3.2.3), where blue colour indicates a CSS and red colour an EBP for a migration rate *m* = 0.01. The dashed line within the red region indicates where CSS and EBP delineate when *m* = 0.1. The gray region indicates parameter values where 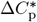 is positive and hence not biologically feasible. **A)** In this panel we have assumed that 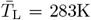 and 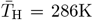.**B)** In this panel we have assumed that 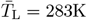 and 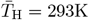.

##### Population initially adapted to habitat L

In this biological scenario we suppose that the population initially inhabited habitat L by assuming that 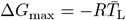 ln *K*^opt^ and 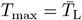.In this case, the convergent stable singularity (eq. 19) simplifies to 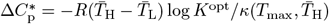 which is always positive (i.e., it lies in the gray region in Figure 9). We thus conclude that in this scenario Δ*C*_p_ evolves to the boundary of the feasible set (i.e. towards 0). Because this boundary is not a singularity further evolution through evolutionary branching is not possible.

##### Initially population is adapted to habitat H

In this biological scenario we suppose that the population initially inhabited habitat H by assuming that 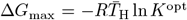 and 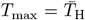.In this case, the convergent stable singularity (eq. 19) simplifies to

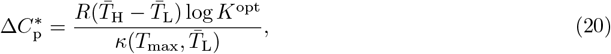

which is always negative and hence biologically feasible (i.e., it lies in the blue region in Figure 9). This singularity is always a CSS because the ESS-condition 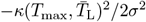 is always negative.

We thus conclude that if the population was initially adapted to either of the habitats, and mutations target only Δ*C*_p_, the heat capacity will evolve either towards 0 (if initially adapted to L), or to 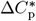 as given in eq. (20) (if initially adapted to H). In both situations no further evolution is expected and hence polymorphism in Δ*C*_p_ is not expected to arise in these two biological scenarios.

##### Large thermal fluctuations

Here, we briefly explore the role of thermal fluctuations on Δ*C*_p_ in the two biological scenario discussed above (numerical results, not shown). First, similarly to above, the singularity is positive and hence not biologically feasible if the population is initially adapted to L. Second, in the alternate scenario where population is initially adapted to H, we find a unique convergent stable singularity that depends on thermal fluctuations. While the full expression is too lengthy to analyse, our numerical investigation indicates that increasing thermal fluctuations lead to an increase in the singular value. Moreover, for the parameter values we tested, the singularity is always a CSS. In this biological scenario, thermal fluctuations thus favour wider CSS protein stability curves.

#### 3.2.4 Co-evolution of thermodynamical parameters under biophysical trade-offs

In this analysis, we assume that mutations can affect multiple thermodynamic parameters simultaneously. Since mutations are likely to influence these parameters in a correlated manner, we introduce trade-offs between them (Mylius and Diekmann, 1995; Rueffler et al., 2004). We focus on the co-evolution of two thermodynamic parameters at a time, without presupposing a specific form for the (potentially empirically unknown) trade-off curve. Instead, we classify trade-off geometries based on their impact on evolutionary dynamics, specifically whether they lead to local (CSS) or global (EBP) adaptation of protein stability. Our methodology follows the approach outlined by Kisdi (2006, see summary on pg. 961-963), which builds on earlier works by de Mazancourt and Dieckmann (2004); Rueffler et al. (2004); Bowers et al. (2005). This method extends the framework of Levins (1962) to incorporate frequency-dependent selection.

For each pair of parameters, we numerically summarize our results by depicting 9 different sets of trade-off curves (e.g., see Figure 10 on the evolution of *T*_max_ and Δ*G*_max_). Each set is constructed (i) according to which singular point they intersect (see the 9 equally spaced black dots in Figure 10A), and (ii) according to the curvature of the trade-off curve at the singularity (indicated by green and black curves intersecting each singularity). Each set of trade-off curves is thus divided into three subsets: first, a subset of trade-off curves whose curvature is greater than that indicated by the green curve, causing the singularity to be a CSS; second, a subset of trade-off curves whose curvature is greater than that indicated by the black curve, causing the singularity to be a repellor; and third, a subset of trade-off curves whose curvature lies between the green and black curves, indicated by the gray region, causing the singularity to be an EBP (see the three light gray trade-off curves intersecting the singularity 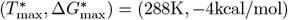 in Figure 10A).

**Figure 10:**
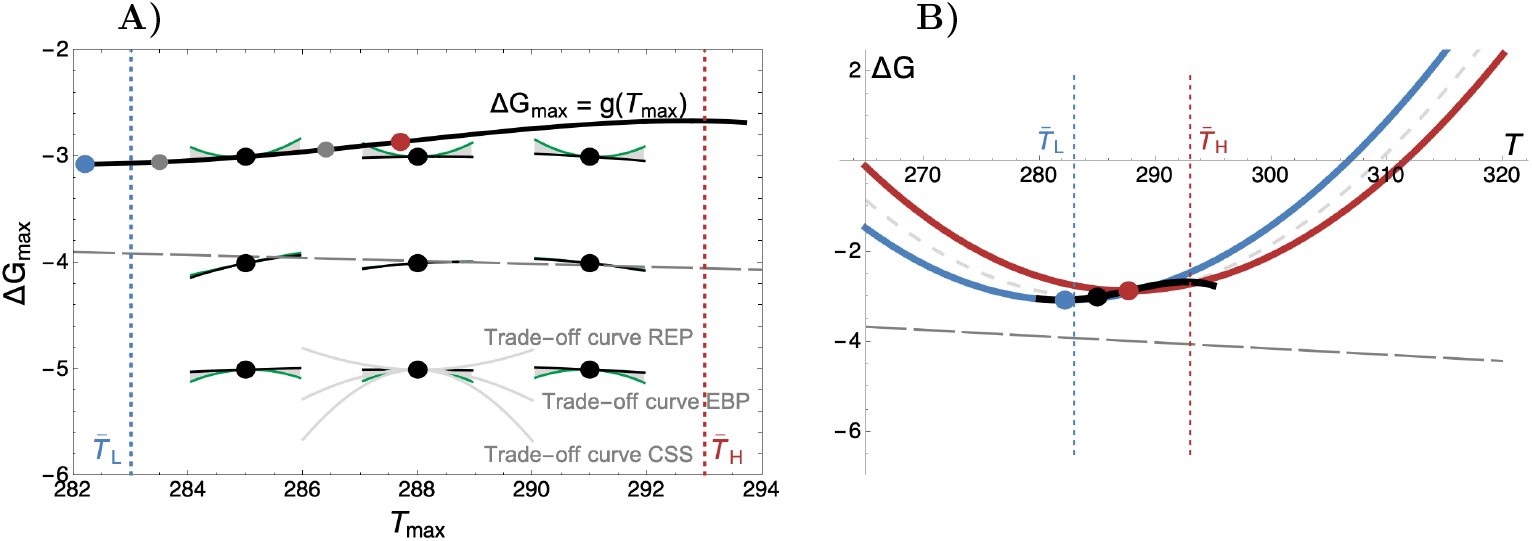
**A)** Classification of singular 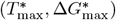 subjected to a trade-off between *T*_max_ and Δ*G*_max_. We plot 9 singular points (black dots), and divide their neighborhood into three regions according to the evolutionary property of the singularity. If the curvature of the trade-off is greater than the curvature of the green curve, then the singularity is an CSS (the bottom gray trade-off curve intersecting the singularity 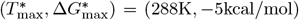 results in a CSS), if the curvature of the trade-off is greater than the curvature of the black curve, then the singularity is a repellor REP (the top gray trade-off curve results in a REP), and if the curvature of the trade-off is in-between black and green curves, then the singularity is an EBP (the middle gray trade-off curve results in an EBP). As an example, we depicted a trade-off curve intersecting the singularity 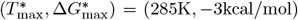,and since its curvature at the singularity lies in the gray region in-between green and black curves, the singularity is an EBP (black curve). This implies that local adaptation via gradual evolution is possible, and we depict the polymorphic CSS by the red and blue dots. The grey dots indicate potential intermediate mutations along the trade-off function, and in panel B we plot the locally adapted adapted protein stability curves corresponding to the red and blue dots. The parameter values are 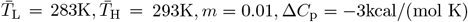.**B)** Two locally adapted protein stability curves (red and blue curves) subjected to a trade-off (black curve). The red and blue curves and dots correspond to the red and blue dots in panel A. The gray dashed curve is the stability curve at the singular value (black dot).

##### Trade-off between Δ*G*_max_ and *T*_max_

Here, we consider a trade-off between Δ*G*_max_ and *T*_max_ expressed as Δ*G*_max_ = *g*(*T*_max_). Figure 10 illustrates our findings on the adaptive process, with fixed parameters 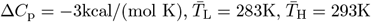, and *m* = 0.01. We verify in Supplementary material S1.3 that varying these parameter values does not qualitatively alter the results (see further discussion below and Supplementary material S1.3 and S2 for details).

The results show a general trend where trade-off curves with strong curvature away from the optimal stability (Δ*G*^opt^) tend to promote global adaptation (CSS), whereas those with moderate curvature away from the optimal stability encourage local adaptation (EBP). Curvatures toward the optimal stability typically lead to a repellor (REP) at the singularity. This pattern arises because strong curvatures away from optimal stability favor intermediate values of *T*_max_, where Δ*G*_max_, and consequently protein stability, are closest to the optimal value. Specifically, any deviation from 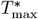 results in a less favorable shift in the protein stability curve, moving it further from Δ*G*^opt^ and making such mutations less likely to be selected. Trade-off curves that open up away from the optimal stability thus lead to a singularity that is a CSS, promoting global adaptation. Conversely, when the trade-off curve points towards the optimal stability, 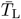 or 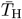 are favored as Δ*G*_max_ shifts towards Δ*G*^opt^, rendering intermediate *T*_max_ a repellor and hindering local adaptation through gradual change. However, trade-off curvatures that balance these two effects lead to a singularity that is an EBP, supporting local adaptation through gradual evolution. Local adaptation is further favoured for narrower protein stability curves, i.e., for smaller values of Δ*C*_p_, and, greater is the difference between the mean 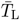 and 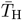 in the two habitats.

##### Trade-off between Δ*C*_p_ and *T*_max_

In this analysis, we examine a trade-off between Δ*C*_p_ and *T*_max_ defined by the function Δ*C*_p_ = *g*(*T*_max_). Note that the function *g* has been redefined for this context. To understand the joint evolutionary dynamics of Δ*C*_p_ and *T*_max_, we differentiate between scenarios where the protein stability curve at temperatures 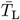 and 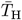 lies either above or below the optimal stability Δ*G*^opt^. In cases where the protein stability curve lies above Δ*G*^opt^, strongly negative curvatures in the trade-off function tend to favor CSS, while strongly positive curvatures favor CSS when the curve lies below Δ*G*^opt^ (as shown in Figure 11, panels A and B, respectively). This difference arises because Δ*C*_p_ affects the width of the protein stability curve. As a consequence, in scenarios where the stability curve lies above Δ*G*^opt^, increasing Δ*C*_p_ (broadening the curve) enhances the likelihood of survival in both habitats. Conversely, when the curve lies at 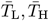 below Δ*G*^opt^, decreasing Δ*C*_p_ (narrowing the curve) enhances survival. Therefore, if the stability curve is above Δ*G*^opt^ and the curvature is strongly negative, mutations that move *T*_max_ closer to 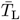 or 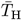 simultaneously narrow the stability curve, creating a disadvantage in the opposite habitat. This leads to global adaptation and favors intermediate *T*_max_ values. On the other hand, positive curvature, which causes wider stability curves, result in mutations towards 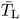 that not only benefit habitat L but also habitat H, due to the curve’s expansion, leading to bistability and making the singularity a REP. Curvatures that balance these opposing effects result in an EBP. If the stability curve lies at 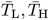 below Δ*G*^opt^, the opposite pattern holds. These dynamics are illustrated in Figure 11, where panels A and B depict scenarios of the protein stability curve being above and below the optimal stability, respectively.

**Figure 11:**
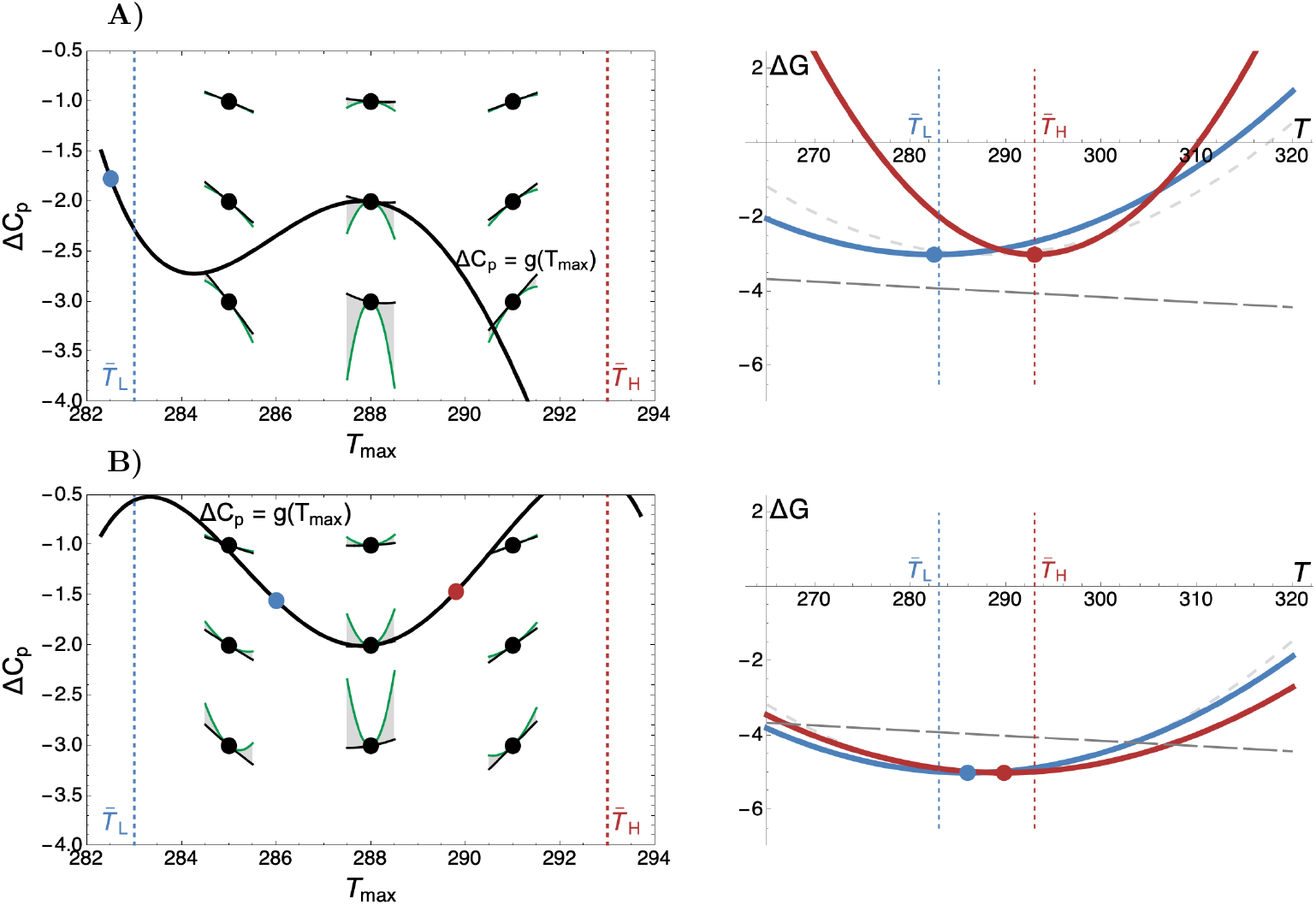
Classification of singular 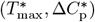 (black dots) subjected to a trade-off between *T*_max_ and Δ*C*_p_. **A)** Assuming Δ*G*_max_ = − 3kcal/mol, we have depicted a trade-off curve intersecting a singularity 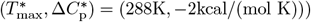 that is an EBP, implying local adaptation via gradual evolution. Two locally adapted stability curves corresponding to the polymorphic CSS (blue and red dots) are depicted in the right panel **B)** Assuming Δ*G*_max_ = − 5kcal/mol, we have depicted a trade-off curve intersecting a singularity 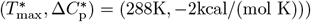 that is an EBP, implying local adaptation via gradual evolution. Two locally adapted stability curves corresponding to the polymorphic CSS (blue and red dots) are depicted in the right panel. The remaining parameter values in both panels are 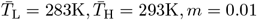.

##### Trade-off between Δ*G*_max_ and Δ*C*_p_

In this analysis, we examine the trade-off between Δ*G*_max_ and Δ*C*_p_, described by the function Δ*G*_max_ = *g*(Δ*C*_p_). Note that the function *g* has again been redefined for this context. As with previous analyses, we differentiate between scenarios where the protein stability curve at temperatures 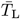 and 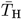 lies either above or below the optimal stability Δ*G*^opt^. This distinction is visually represented by the blue and red horizontal lines in Figure 12. Additionally, we consider two cases: one where 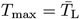 (panels A) and another where 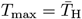 (panels B).

**Figure 12:**
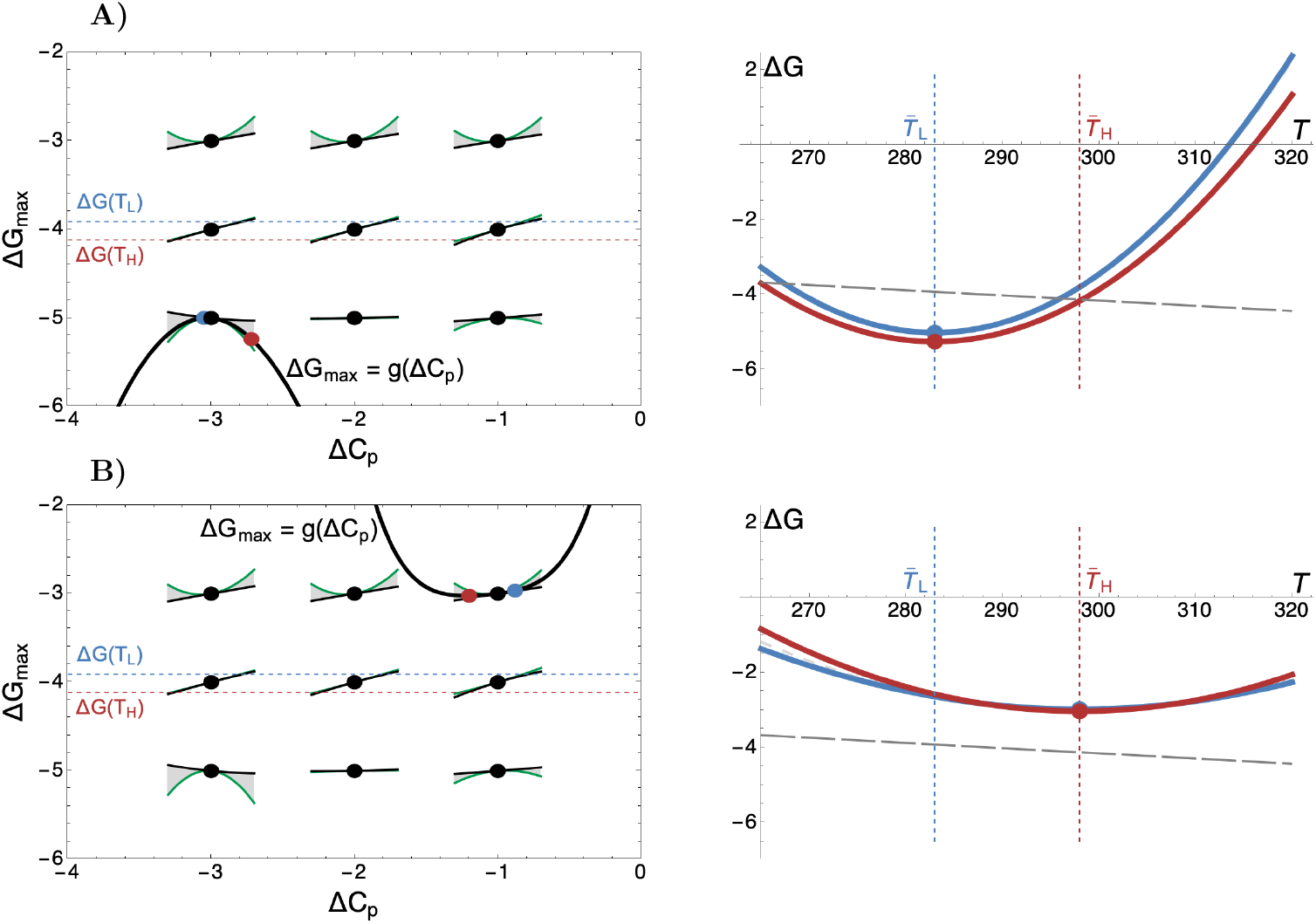
Classification of singular 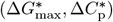 subjected to a trade-off between Δ*G*_max_ and Δ*C*_p_. **A)** Assuming 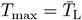,we have depicted a trade-off curve intersecting a singularity 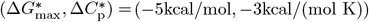 that is an EBP, implying local adaptation via gradual evolution. Two locally adapted stability curves corresponding to the polymorphic CSS (blue and red dots) are depicted in the right panel **B)** Assuming 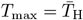,we have depicted a trade-off curve intersecting a singularity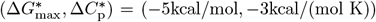 that is an EBP implying local adaptation via gradual evolution. Two locally adapted stability curves corresponding to the polymorphic CSS (blue and red dots) are depicted in the right panel. The remaining parameter values in both panels are 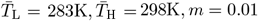.

In panels A, a trade-off curve with a negative curvature is depicted, which leads to a singularity that is an EBP. This allows for the possibility of local adaptation through gradual evolution. Stronger negative curvatures favor a CSS. The reasoning behind this pattern is analogous to that discussed for trade-offs between Δ*G*_max_ and *T*_max_. A singularity is favored by selection and becomes a CSS when Δ*G*_max_ is closest to Δ*G*^opt^ at the singularity. If Δ*G*_max_ is furthest from Δ*G*^opt^ at the singularity, with trade-off curves pointing towards the line of optimal stability, the singularity becomes a repellor (REP). This pattern is observed consistently in both cases (as shown in panels A and B).

#### 3.2.5 Linear model of protein stability

Here, similarly to the homogeneous population model, we analyse the evolution of *T*_m_. We find a unique convergent stable singularity,

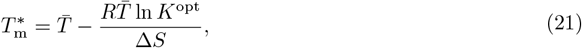

where 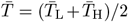 is the average temperature across habitats. Note that the singularity (eq. 21) has a similar form to that in the homogeneous population case (eq. 14), but where the mean temperature of a single habitat is replaced by the average temperature over multiple habitats. As in the homogeneous population case, the singularity (eq. 21) is independent from thermal fluctuations. Furthermore, the singularity (eq. 21) is a CSS when

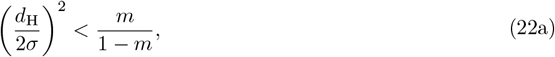

where

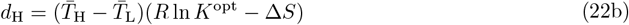

is the sum of the distances between the optimum stability and the protein stability in each habitat, that is, 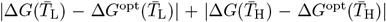.Evolutionary branching is favoured when this distance increases, as selection against maladapted alleles strengthens with increasing distance. Moreover, evolutionary branching is favoured when *σ* decreases, *m* decreases, while fluctuations *ϵ* have no effect.

## 4 Discussion

This study explored the adaptive evolution of protein thermodynamic stability in complex thermal environments by employing a mathematical model that integrates two previously established components: a population genetic model representing species distributed across thermally distinct habitats, and a thermodynamic model of protein stability that incorporates temperature-dependent enthalpy Δ*H*(*T*) and entropy Δ*S*(*T*) contributions. For comparison, we also analyzed a simplified model with temperature-independent enthalpy and entropy. To our knowledge, no previous work has examined the evolutionary dynamics of protein stability in the context of complex thermal environments or by incorporating temperature-dependent enthalpy and entropy contributions.

In the simplified model, where Δ*H* and Δ*S* are independent of temperature, protein stability is linear with respect to temperature. The adaptive process is primarily driven by the mean temperature of the thermal environment because, in the linear model, thermal fluctuations around the mean cancel out. Nonetheless, local adaptation of protein stability can still occur if mean temperatures differ sufficiently across habitats, leading to divergence in the melting temperature *T*_m_. However, while the linear model adequately describes protein stability near the melting temperature, it fails to capture deviations observed in more general thermal environments. The model with temperature-dependent enthalpy and entropy contributions, which render protein stability non-linear, addresses these limitations by introducing three key thermodynamic parameters: maximum stability Δ*G*_max_, the temperature of maximum stability *T*_max_, and the heat capacity change Δ*C*_p_. These parameters define the non-linear stability profile and result in more complex adaptive responses across diverse thermal conditions. Below, we present our findings and, where relevant, compare them to predictions from the linear model.

### Small shifts in thermal environments

We first examined a population in a single habitat and the selective pressures imposed by small shifts in the mean temperature. Our analysis shows that changes in mean temperature 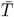 can lead to distinct adaptive responses. When 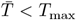, an increase in mean temperature 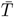 tends to favor mutations that elevate *T*_max_, while also promoting increases in Δ*G*_max_ and decreases in Δ*C*_p_. Conversely, when 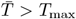, the opposite pattern occurs. This reveals that thermal adaptation depends on the evolutionary history of the protein, as seen in other contexts in previous studies on protein evolution (Shah et al., 2015; Starr and Thornton, 2016).

This finding highlights a crucial difference between the linear and non-linear models of protein stability. In the linear model, increasing mean temperature selects for higher melting temperatures *T*_m_, but in non-linear models, the adaptive response can push proteins toward lower *T*_m_, depending on the relationship between *T*_max_ and 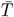.This suggests that thermal adaptation is more nuanced than predicted using linear models, with proteins adopting different stability profiles based on their evolutionary context.

We next examined how thermal fluctuations impact protein stability. Our findings indicate that Δ*G*_max_ and *T*_max_ are relatively insensitive to fluctuations (but see below the effect of *T*_max_ on local adaptation). In contrast, Δ*C*_p_, which controls the width of the stability curve, is highly sensitive to fluctuations. Proteins with broader stability curves (higher Δ*C*_p_) are better suited to environments with significant temperature variability, as their stability spans a wider range of temperatures. This indicates that while mean temperature determines core stability features, fluctuations exert strong selective pressure on the protein’s ability to withstand variable conditions.

### Local and global adaptation

To investigate patterns of local and global adaptation, we modeled populations inhabiting two thermally distinct habitats, connected by migration. Our results show that local adaptation of protein stability is expected when the difference between the mean temperatures of the two habitats is large enough. Specifically, the thermodynamic parameters Δ*G*_max_ and *T*_max_ are predicted to undergo local adaptation in these cases, reflecting distinct stability profiles for each habitat. This is consistent with empirical studies, which suggest that Δ*G*_max_ is a primary driver of differentiation between cold- and warm-adapted proteins (Sawle and Ghosh, 2011; Pucci and Rooman, 2014).

In contrast, Δ*C*_p_ does not appear to undergo local adaptation. Comparative studies support this finding, with evidence showing that Δ*C*_p_ is less differentiated between habitats compared to Δ*G*_max_ (Graziano, 2008; Ragone, 2004). Proteins with narrower stability curves, indicated by lower Δ*C*_p_, tend to exhibit stronger local adaptation in Δ*G*_max_ or *T*_max_, as narrower curves impose higher fitness costs in the alternate habitat. This suggests that proteins adapted to local environments will exhibit narrower stability curves compared to globally adapted proteins, a hypothesis that warrants future empirical testing.

### Thermal fluctuations and local adaptation

Large thermal fluctuations in two connected habitats can result in overlapping temperature ranges, and thus might be expected to hinder adaptation to local environments. However, our findings indicate the opposite: increased fluctuations actually promote local adaptation of protein stability. This counter-intuitive result arises because thermal fluctuations increase the occurrence of extreme temperatures, which impose higher fitness costs due to the non-linear relationship between protein stability and temperature. At these extreme temperatures, protein stability deviates strongly from its optimal range, favouring selection for specialized stability profiles in each habitat. Consequently, selection favors adaptations in *T*_max_ that are tailored to the specific thermal extremes of each environment, while maximum stability Δ*G*_max_ is only mildly affected by fluctuations, and heat capacity change Δ*C*_p_ remains unaffected. This finding underscores the importance of considering both mean temperature and thermal fluctuations when studying protein evolution in heterogeneous environments. Moreover, we expect these results to hold true beyond the context of protein evolution, as similar dynamics may arise in other systems where fitness depends on the interplay between environmental fluctuations and non-linear trait-environment relationships.

### Trade-offs between thermodynamic parameters and local adaptation

Mutations are likely to affect more than one thermodynamic parameter simultaneously, prompting us to investigate the role of trade-offs between these parameters. We identified two main trade-off patterns: one between maximum stability Δ*G*_max_ and either maximum-stability temperature *T*_max_ or heat capacity Δ*C*_p_, and another between Δ*C*_p_ and *T*_max_. Our analysis showed that for the first case, trade-off curves where Δ*G*_max_ is closest to the line of optimal stability for intermediate temperatures 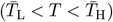 tend to favor global adaptation. Specifically, if Δ*G*_max_ *>* Δ*G*^opt^(*T*), convex trade-offs favor global adaptation, whereas if Δ*G*_max_ *<* Δ*G*^opt^(*T*), concave trade-offs favor global adaptation. This pattern arises because deviations from optimal stability is smallest for intermediate temperatures, thus promoting global adaptation. Conversely, the opposite pattern favors bi-stability, where locally adapted protein stability profiles can only be reached if mutational effects are large. A balance between these effects results in gradually evolving local adaptation.

In the case of trade-offs between Δ*C*_p_ and *T*_max_, we observe that when Δ*G*_max_ *>* Δ*G*^opt^(*T*) for 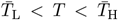,concave trade-offs favor global adaptation. This occurs because Δ*C*_p_ is the largest (least negative) for intermediate *T*_max_ values, making the protein stability curve widest, implying that any minor deviation narrows the protein stability curve, intensifying within-habitat selection in both habitats and favouring global adaptation. The opposite argument can be used for Δ*G*_max_ *<* Δ*G*^opt^(*T*) for 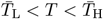.Trade-off curves with opposite curvatures then favour bi-stability, where locally adapted protein stability profiles can only be reached if mutational effects are large, and a balance between these patterns result in an gradual local adaptation. These insights highlight that understanding the nature of trade-off geometries is important in reliably predicting protein stability evolution.

### Greater differentiation in adaptation from colder to warmer habitats

Our results suggest that local adaptation leaves stronger imprints on protein stability when populations from colder habitats adapt to warmer conditions, than in the reverse scenario. Proteins originating from colder habitats are farther from their optimal stability when exposed to higher temperatures, necessitating larger mutational changes for adaptation. This finding aligns with studies on protein thermal adaptation, which indicate that the cost of protein destabilization is higher at elevated temperatures, leading to stronger selective pressure for stability (Chen and Shakhnovich, 2010; Agozzino and Dill, 2018; Berger et al., 2021).

Interestingly, recent empirical studies have found that mutations affecting protein stability rise to higher frequencies more quickly in warmer environments compared to colder ones (Zheng et al., 2024). This supports the idea that protein stability is under stronger selection pressure in warm habitats, where destabilization has more immediate fitness consequences. Our model extends these insights by showing how thermal adaptation to warmer habitats can leave distinct signals in protein stability curves, offering potential applications for reconstructing demographic and evolutionary histories.

### Protein stability as a tool for detecting hidden signals of adaptation

The thermodynamic parameters we studied – Δ*G*_max_, *T*_max_, and Δ*C*_p_ – provide a powerful framework for detecting hidden signals of thermal adaptation. First, they directly link temperature to an evolving phenotype (protein stability), and can be estimated both experimentally and computationally (Fersht, 1999; Barrick, 2018; Thurlkill et al., 2023; Seeliger and De Groot, 2010; Kim et al., 2016; Timr et al., 2020; Galano-Frutos et al., 2023; Wang et al., 2024). Recent advances in computational tools also enable the prediction of how genetic mutations affect thermodynamic stability and, consequently, allelic fitness directly from DNA sequences (Pancotti et al., 2022; Blaabjerg et al., 2023).

Second, these parameters help researchers identify adaptations that might otherwise be overlooked using conventional genotype-phenotype analyses. Protein stability is particularly likely to be a key target for natural selection in systems where phenotypes must be maintained across varying thermal environments, such as in many core cellular processes (Bomblies, 2020, 2022; Bomblies and Peichel, 2022). By combining our model with population genetic data and computational or empirical estimates of protein stability, we provide a robust framework for linking genetic variation directly to adaptive phenotypes. This approach could also be developed into a tool for detecting signatures of thermal adaptation or determining whether temperature has driven a detected selective sweep. Ultimately, this combined framework addresses critical questions about the role of biophysical traits, such as protein stability, in shaping selection signals observed in genome scans.

## Conclusions

Our study presents a comprehensive framework for understanding the adaptive evolution of protein stability in thermal environments. By examining the roles of mean temperature, thermal fluctuations, and habitat heterogeneity, we provide key insights into the molecular mechanisms driving thermal adaptation, emphasizing the need to account for both temperature averages and variability in evolutionary studies. Future research should aim to empirically validate these models and explore how these adaptive dynamics manifest in natural populations, particularly as environmental conditions continue to change.

## Appendices

### A. Mathematical models

#### A.1 Homogeneous and heterogeneous population genetic models

In the main text (eq. 3), we present the general migration model with arbitrary migration patterns and an arbitrary number of subpopulations and alleles. The single-habitat version of this model, with two alleles, *x* and *y*, and symmetric migration (*m*_*XY*_ = *m* for all *X*≠ *Y*), takes the form

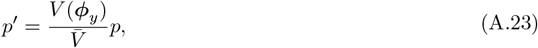

where *p* is the frequency of *y*, and which we refer to as the homogeneous model used in the main text.

The two-habitat and two-allele version with symmetric migration takes the form

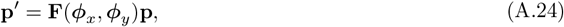

where

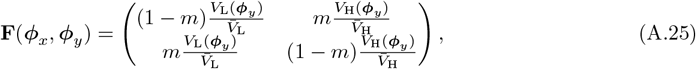

which we refer to as the heterogeneous model used in the main text. The viability fitness functions are provided in the main text in eqs. (1)-(2).

#### A.2 Thermodynamic model of protein stability

The stability of the protein at temperature *T* is defined as the difference in Gibbs free energy of a protein in its folded and unfolded states at *T* (eq. 4). Because enthalphy of a protein in state *X* ∈ {F, U} can be expressed at constant pressure in terms of its heat capacity as d*H*_*X*_ = *C*_p,*X*_ d*T*, and entropy can in turn be expressed as *T* d*S*_*X*_ = *C*_p,*X*_ d*T*, both contributions are dependent on temperature and can be calculated as

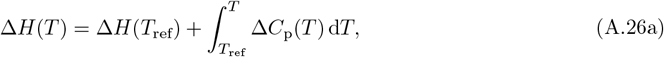

and

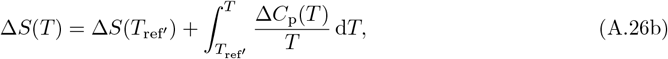

where we used Kirchoff’s law of thermodynamics, and where *T*_ref_, *T*_ref′_ are some arbitrary reference temperatures. For our evolutionary analysis it will be convenient to use *T*_max_ = *T*_ref_ = *T*_ref*′*_as reference temperature, defined as a temperature where the temperature-dependent entropy is 0, that is, Δ*S*(*T*_max_) = 0 (see below for an alternative choice). This choice is practical because then each thermodynamical parameter has a qualitatively distinct effect on the protein stability profile (see main text, Figure 2). That is, the contributions of enthalpy and entropy at *T* are as

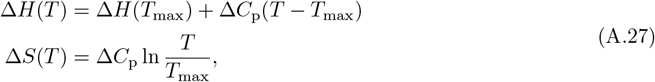

in which case the Gibbs free energy can be written as

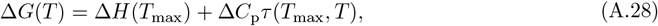

with a temperature-dependent term

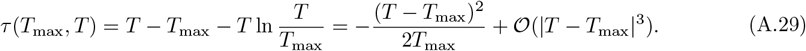

These expressions were derived from eq. (A.26) by assuming that the difference in heat capacity Δ*C*_p_ is independent of temperature, a common assumption in the literature (Becktel and Schellman, 1987; Schellman, 1987; McCrary et al., 1996; Barrick, 2018). In the main text we use *κ*(*T*_max_, *T*) = −(*T* − *T*_max_)^2^*/*(2*T*_max_) as an approximation to *τ* (*T*_max_, *T*), which has also been used previously (Hawley, 1971; Yeritsyan and Badasyan, 2024). Some useful properties of the temperature-dependent term are:

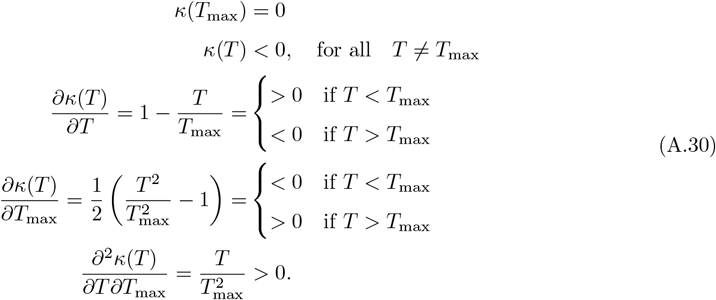

We finally note that elsewhere in literature a common choice for a reference temperature is the melting temperature *T*_m_, at which Δ*H*(*T*_m_) = *T*_m_Δ*S*(*T*_m_), leading to the so-called modified Gibbs-Helmholtz equation 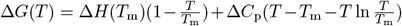 also called a constant heat capacity model (Barrick, 2018). This equation is not be used in this paper.

### B Invasion fitness and weak selection approximation

Because we focus on small mutational effects on the phenotype, we write ***ϕ***_*y*_ = ***ϕ*** + *δ****η***, where ***η*** = (*η*_G_, *η*_T_, *η*_C_) determines the direction of mutation, *δ* is a small parameter that defines the magnitude of the mutational effect, and, for notational simplicity, we denote the phenotype of the resident *x* by ***ϕ*** (in the main text we write ***ϕ***_*x*_). When analyzing the evolutionary dynamics, we always focus on a single thermodynamical parameter at a time by setting two elements of the vector ***η*** to 0, thus effectively studying the evolution of scalar-valued traits. For example, when analyzing the dynamics of Δ*G*_max_, we set *η*_T_ = *η*_C_ = 0.

#### B.1 Homogeneous populations

In a homogeneous population the invasion fitness of a rare *y* in a resident population of *x* is

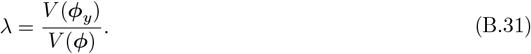

For small *δ* the invasion fitness can be approximated up to second order as

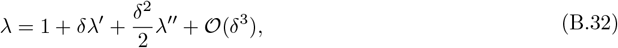

where

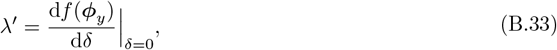

and

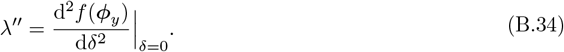

##### First-order term *λ*^*′*^

To calculate the first-order term, we substitute the viability fitness function (eqs. 1-2) into eq. (B.33), and we get

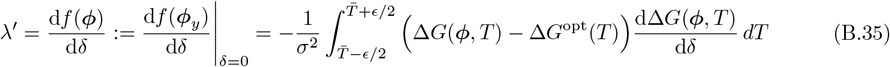

where the derivative inside the integral is

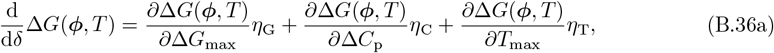

with

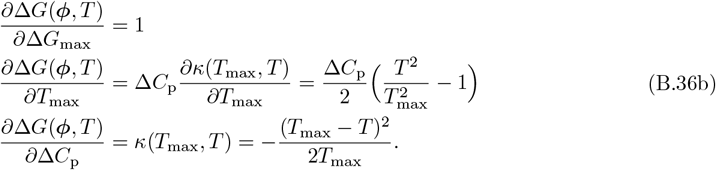

As we analyze the evolutionary dynamics of each parameter separately, we always set the other two elements of the vector ***η*** to 0. Thus, the singular 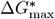 is obtained by solving

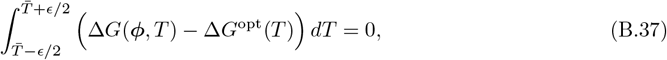

the singular 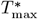 from solving

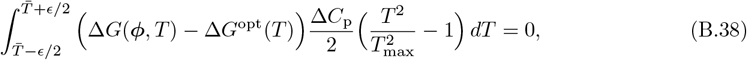

and the singular 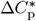 from solving

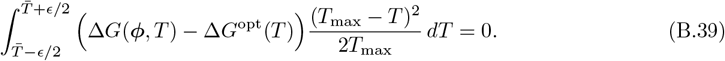

##### Second-order term *λ*^*′′*^

The calculate the second-order term, we substitute the viability fitness function (eqs. 1-2) into eq. (B.34), and we get

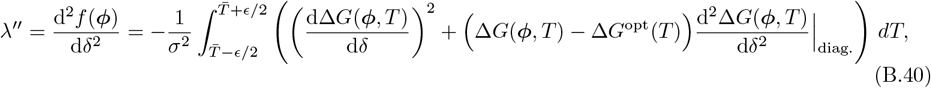

where the first term in the integrand is given in eq. (B.36). Because we analyse each parameter separately, we only consider the diagonal (non-mixed) second-order derivatives

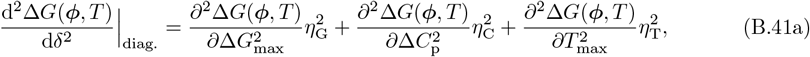

with

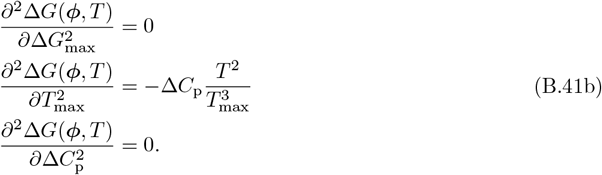

This second-order term (eqs. B.40-B.41) determines whether a convergence stable singularity is a CSS or an EBP, according to eq. (B.52). In Supplementary material S1 and S2, we provide detailed analysis of the evolutionary properties of each evolutionary parameter.

##### Small fluctuations

For small (negligible) thermal fluctuations (*ϵ* = 0), the singularities can be solved from eqs. (B.37)-(B.39) by ignoring the integrals. We will explore these solutions in greater detail in the Supplementary material S1-S2, here we just note that the singular solutions can be divided into solutions where either 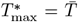 (obtained from the second term in eq. B.38) or where 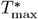 is solved from Δ*G*(0) − Δ*G*^opt^ = 0, which is a quadratic equation in *T*_max_ and hence has at most two solutions. If 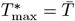, then Δ*C*_p_ is neutral and every value is 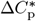. This is because 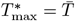 solves eq. (B.39) and hence for every Δ*C*_p_ the equality in eq. (B.39) is satisfied. If 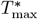 is solved from Δ*G*(0)−Δ*G*^opt^ = 0, then at 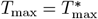, every parameter combination (Δ*G*_max_, Δ*C*_p_) becomes singular since they by default also satisfy Δ*G*(0) − Δ*G*^opt^ = 0 and thus solve eqs. (B.37) and (B.39). The same is true also for 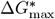 and 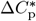 when solved for singularities, as the other two parameters then by default also satisfy the singular solutions.

#### B.2 Heterogeneous populations

In heterogeneous populations, the invasion fitness *λ* of a rare *y* in a resident population of *x* is the dominant eigenvalue of the matrix **F** (eq. A.25) evaluated at **p** = (*p*_L_, *p*_H_) = (0, 0). This matrix takes the form

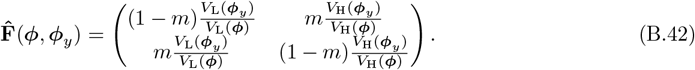

For small *δ*, the invasion fitness up to second order can be approximated as

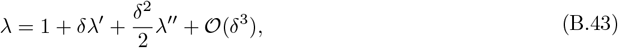

where

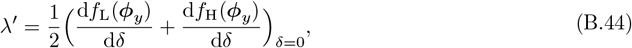

and

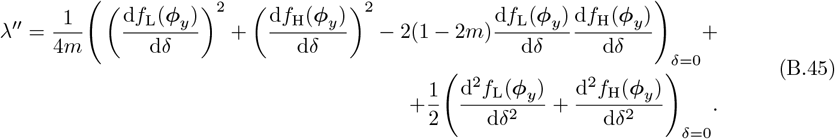

Note that whereas the second-order condition depends on *m*, the first-order condition is independent of migration. These expressions were obtained using Mathematica Wolfram Version 14.0, but can also be calculated analytically (Priklopil, 2025).

##### First-order term *λ*^*′*^

By substituting the viability fitness function (eqs. 1-2) into eq. (B.44), the terms in the brackets (eq. B.44) can be expressed as

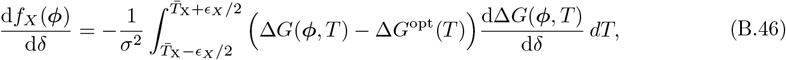

for *X* ∈ {L, H}, where the derivative inside the integral is given as in eq. (B.36). As we analyze the evolutionary dynamics of each parameter separately, we always set the other two elements of the vector ***η*** to 0. Thus, the singular 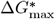 is obtained by solving

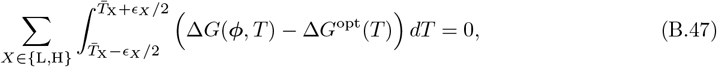

the singular 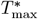 from solving

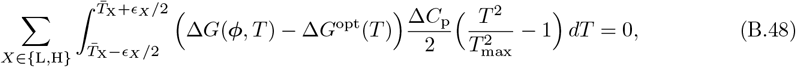

and the singular 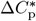 from solving

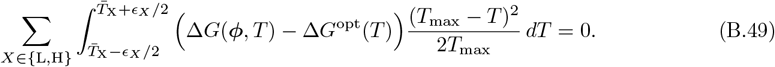

The second-order term (eq. B.45) can be analyzed using eq. (B.46), along with the analogue of eq. (B.40) in combination with eq. (B.41), constructed for heterogeneous populations. The above model with thermal fluctuations is analysed numerically using Mathematica Wolfram Version 14.0, as presented in the main text. Further details can be found in Supplementary material S1-S2.

##### Small fluctuations

For small (negligible) thermal fluctuations (*ϵ* = 0), the singularities can be solved from eqs. (B.47)-(B.49) by ignoring the integrals.

#### B.3 Adaptive evolution and the classification of singularities

Singular evolutionary thermodynamic parameters can be calculated, and their class can be determined by analysing the invasion fitness (Geritz et al., 1998; Rousset, 2004). Supposing that mutations only affect a single evolutionary parameter at a time (by setting two elements of the vector ***η*** to 0), the phenotype ***ϕ***^*^ is a CSS whenever it satisfies

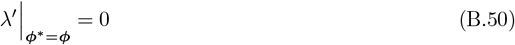

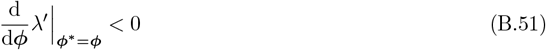

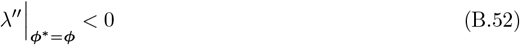

(Geritz et al., 1998; Della Rossa et al., 2015). Singularity ***ϕ***^*^ is solved from the first equality (eq. B.50), the convergence stability of this singularity is determined from the second inequality (eq. B.51), and the final inequality (eq. B.52) guarantees that the singularity is a CSS. If the final inequality has an opposite sign the singular ***ϕ***^*^ is an evolutionary branching point (EBP). Note also that for the homogeneous model, the convergence stability-condition (eq. B.51) and the ESS condition (eq. B.52) are in fact the same (compare with eqs. B.33-B.34). A case where mutations affect multiple parameters simultaneously is analysed in Supplementary material S1.3.1 and S2.

## S1 Supplementary material: mathematical analysis

### S1.1 Homogeneous populations

Here we provide details on the evolutionary dynamics of thermodynamic parameters in the homogeneous population model. Further analysis and numerical explorations are provided in Supplementary material S2.

#### S1.1.1 Temperature of maximum stability

##### Small thermal fluctuations

For small thermal fluctuations (*ϵ* = 0) the singular temperature of maximum stability can be solved from eq. (B.38) by ignoring the integral, and we find one (biologically unrealistic) negative solution 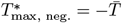 and three positive solutions

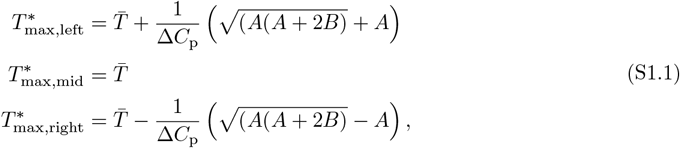

where

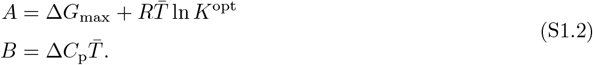

Note that in the main text we used a different parametrization, but we use the above notation here because it simplifies the calculations (see below). Because for biologically realistic values *B* is negative and large (of order 𝒪(10^2^)), we find that that the solutions 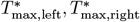 are real solutions (the discriminant is positive) whenever *A* ≤ 0, that is, whenever 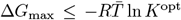. Whenever 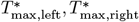 are real they thus lie on both sides of the mid-solution 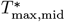, and note that when 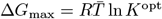, the two solutions coincide with 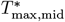. Moreover, we find that the CSS condition for the mid-solution 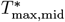 is

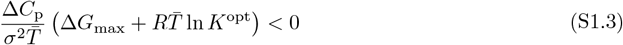

which is satisfied whenever 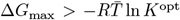. This implies that for 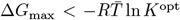 the mid-solution 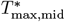 is a repellor and 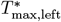 and 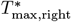 are CSSs.

Model parameters have no effect on the mid-solution 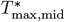 (except that, trivially, increasing 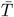 will increase 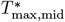). The effect of parameters on the left 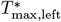 and right solutions 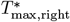 is slightly more involved. The effect of Δ*G*_max_ can be found from

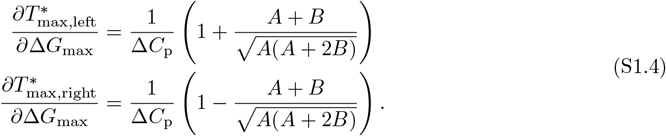

Because *B <* 0 and whenever 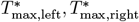 are real then *A <* 0, implying that the fraction within the brackets is negative. From this immediately follows that the second partial derivative is always negative and 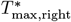 decreases with increasing Δ*G*_max_. It is also simple to show that 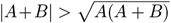 and hence the fraction inside the brackets is always smaller than −1, implying that the first partial derivative is always negative and 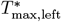 increases with increasing Δ*G*_max_. The effect of Δ*C*_p_ can be found from

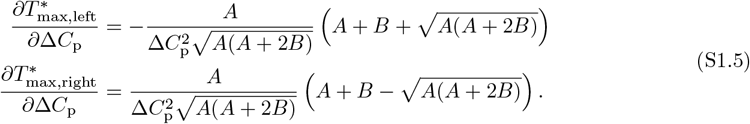

Because 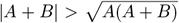 and *A, B <* 0, we find that with increasing Δ*C*_p_ the solution 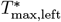 decreases and 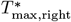 increases.

The effect of *K*^opt^ can be found from

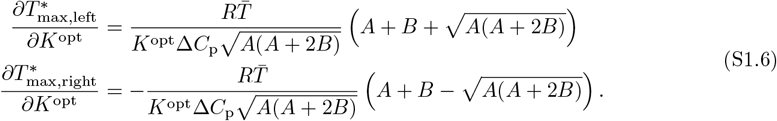

Using the same logic as above, we find that as *K*^opt^ increases, the solution 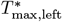 increases and 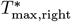 decreases. Finally, the effect of 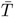 can be found from

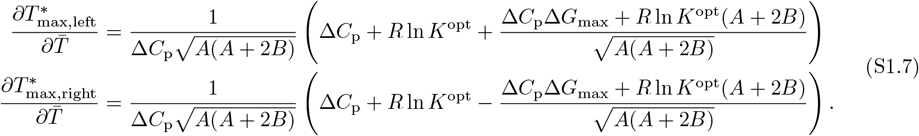

For biologically realistic values our numerical exploration indicates that with increasing 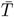 both solutions 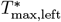 and 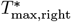 increase.

##### Large thermal fluctuations

Let us now investigate the effect of thermal fluctuations that are not negligible (0 *<< ϵ*). To find the singularities we must solve eq. (B.38), which is algebraically challenging. We can obtain the algebraic expressions using Mathematica, but the algebraic expressions are legnthy and are not presented here. Nevertheless, we find that solving eq. (B.38) is equivalent to solving a quadratic polynomial in 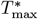, implying there exist at most four solutions. Moreover, our numerical analysis suggests that the behavior of solutions is qualitatively as well as quantitatively similar to the analytical analysis of solutions with small fluctuations as presented above. That is, first, one out of four solutions is always negative and hence not biologically relevant. Second, below certain threshold of Δ*G*_max_, the three positive solutions (left, middle, right) are real valued, with the middle solution being always a repellor and the left and right solutions are always CSSs (see Figure 13). Third, at the threshold, the left and the middle solutions coalesce and become a pair of complex conjugate solutions above the threshold. Above the threshold this pair of solutions is thus not biologically relevant. Finally, the right solution remains to be real-valued CSS for all Δ*G*_max_ (see Figure 13 for illustration). In Figure 14 we plot the effect of Δ*C*_p_ and Δ*G*_max_ on the singular solutions, which is also closely similar to the case of small fluctuations.

**Figure 13:**
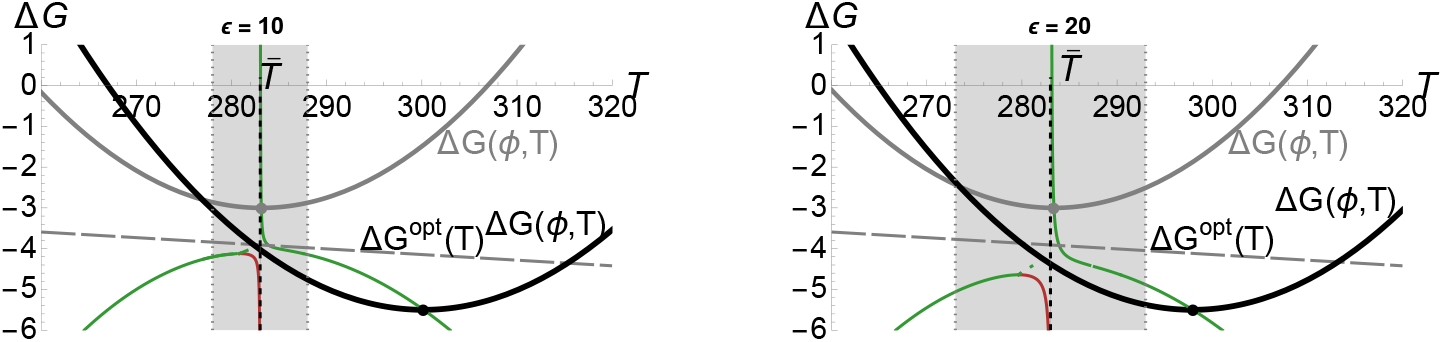
Singular solutions 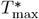 for a homogeneous model with thermal fluctuations. We use the same labeling as in the main text, with gray region indicating the range of temperatures the individuals experience in the homogeneous population model.

**Figure 14:**
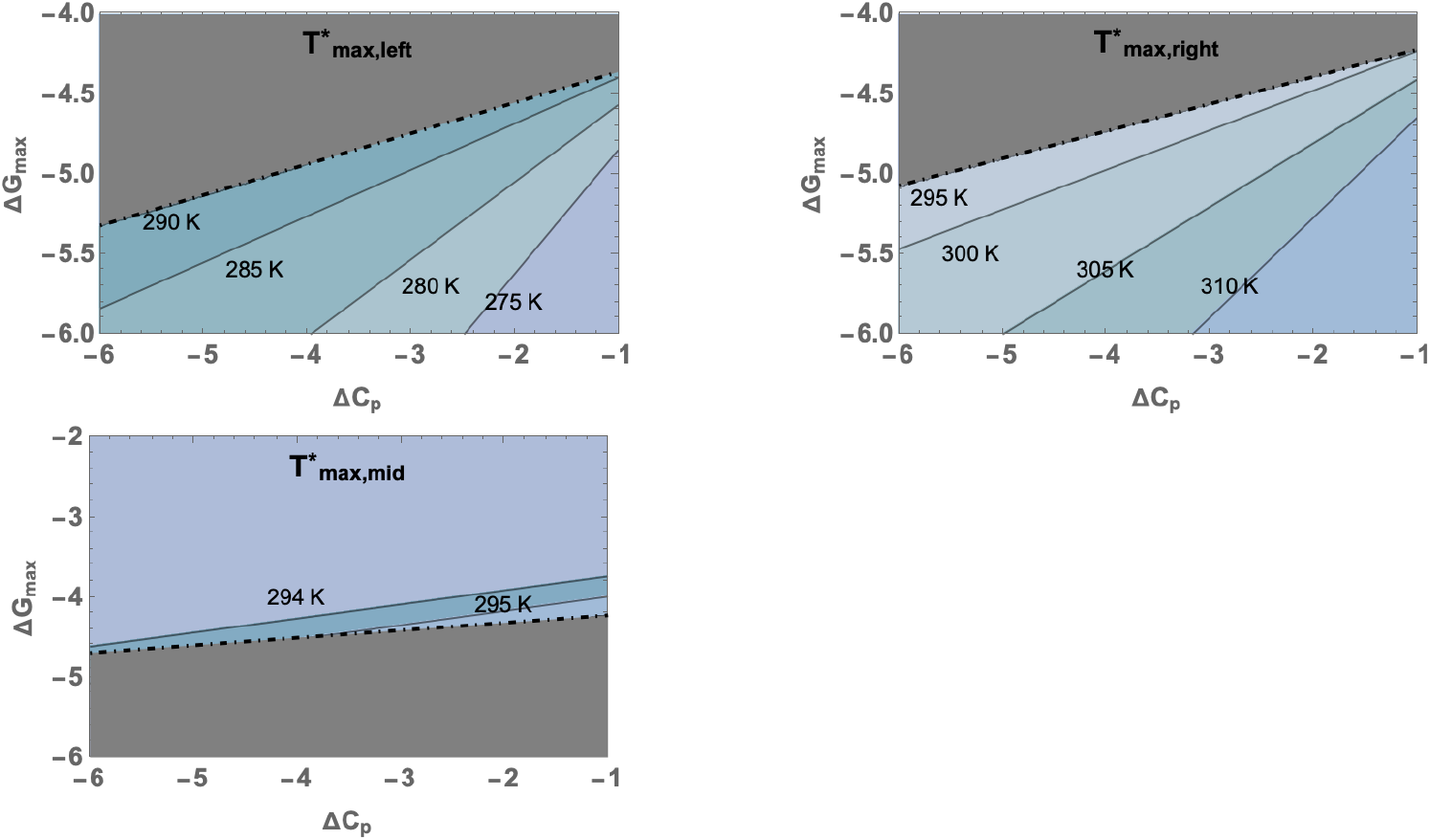
Contourlines for singular 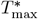 as a function of (Δ*C*_p_, Δ*G*_max_). The gray region indicates the parameter values for which the corresponding 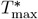 is complex and hence not biologically relevant.

**Figure 15:**
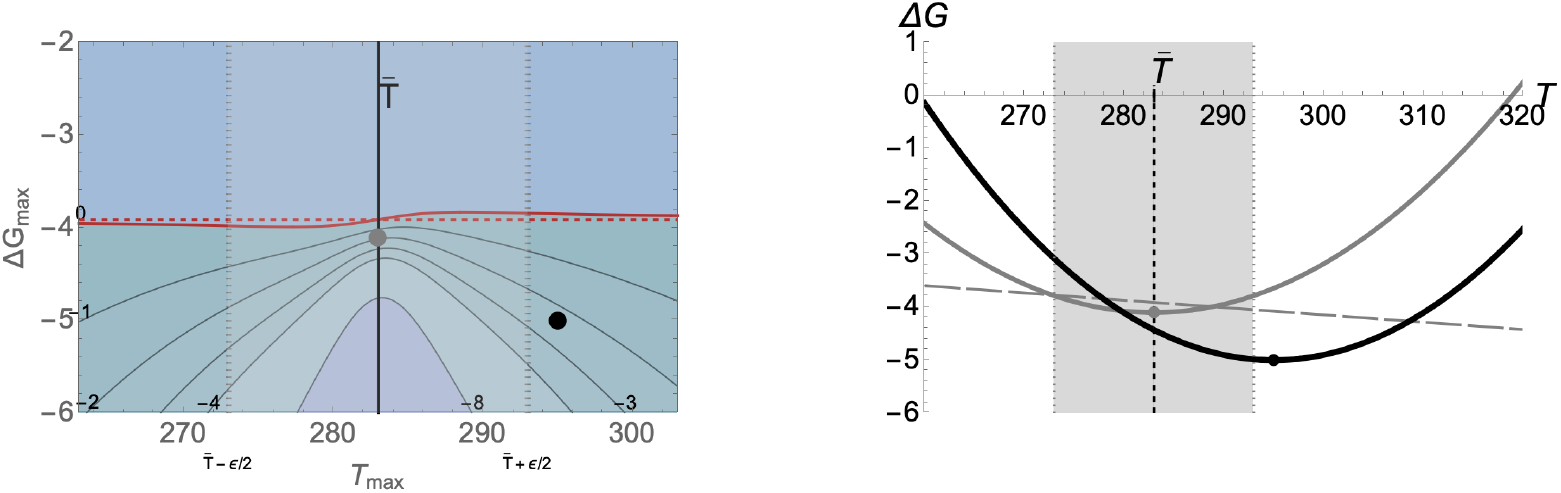
**Left panel:** Contourlines for singular 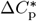 as a function of (*T*_max_, Δ*G*_max_). The red curve indicates the critical threshold (eq. S1.11) above which Δ*C*_p_ is positive and hence not biologically feasible. As *ϵ* approaches 0, this critical threshold approaches the dashed red-line where 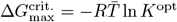. The gray and black dots indicate the parameter combinations used in the right panel. **Right panel:** Protein stability curves for CSS 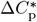 calculated for two parameter combinations. The gray protein stability curve is obtained using (Δ*G*_max_, *T*_max_) = (−4.1kcal/(mol K), 293K) as indicated with a gray dot, which yields a CSS 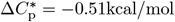 and which corresponds to the gray dot in the right panel. The black curve is obtained using (Δ*G*_max_, *T*_max_) = (−4.5kcal/(mol K), 305K) as indicated with a black dot, yielding 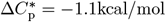 which corresponds to the back circle in the right panel.

#### S1.1.2 Heat capacity

##### Small thermal fluctuations

For small thermal fluctuations (*ϵ* = 0) the singular heat capacity is a a CSS because the (convergence stability and ESS) condition

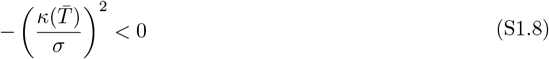

is always satisfied.

##### Large thermal fluctuations

Let us now investigate the effect of thermal fluctuations that are not negligible (0 *<< ϵ*). To find the singularities we must solve eq. (B.39), and we find

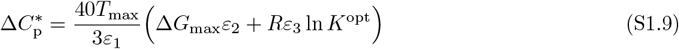

where

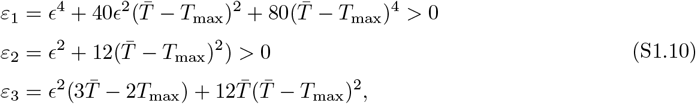

where *ε*_3_ can be negative or positive but for biologically reasonable values we ought to have 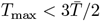 for which always *ε*_3_ *>* 0 (because 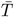 is usually around 300*K*, the maximum stability should not exceed 450*K* which for biologically realistic proteins never does). Note that in the main text we use an alternative parametrization. The singular 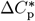 (eq. S1.9) is negative whenever the term in the brackets is negative, and this is true whenever

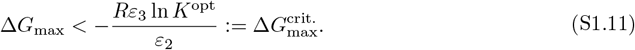

Note that by letting *ϵ* to go to 0, then eq. (S1.9) reduces to eq. (11) and 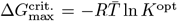. The singular 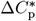 (eq. S1.9) is a CSS whenever

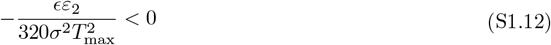

which is always satisfied.

The effects of parameters Δ*G*_max_ and *K*^opt^ on 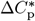 can be found from eq. (S1.9). Because Δ*G*_max_ and *K*^opt^ are multiplied by expressions that are positive, increasing Δ*G*_max_ and *K*^opt^ will increase 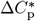. To understand the effects of 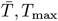, *T*_max_ and *ϵ* is more involved, and we will here only look at the case where 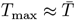. We get

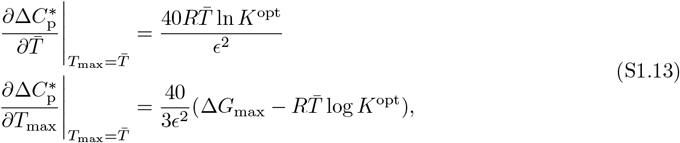

and thus increasing 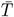 will always increase Δ*C*_p_, and increasing *T*_max_ will always decrease Δ*C*_p_. Finally, the effect of *ϵ* for can be obtained from

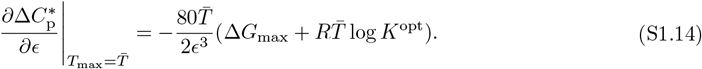

We find that increasing thermal fluctuations *ϵ* will increase Δ*C*_p_ whenever 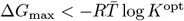 which is approximately satisfied for 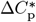 to be negative.

### S1.2 Heterogeneous populations

Here we provide details on the evolutionary dynamics of thermodynamic parameters in the heterogeneous population model. Further analysis and numerical explorations are provided in Supplementary material S2.

#### S1.2.1 The effect of parameters on the singular maximum-stability temperature

For negligible thermal fluctuations (*ϵ* = 0), but otherwise general biological scenarios, the first-order term (eqs. B.44, B.46, B.49) becomes

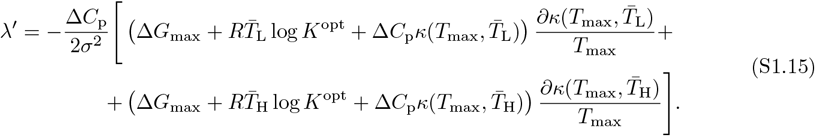

The solutions are lengthy and difficult to interpret, but they simplify considerably if we assume that the population initially inhabited and was adapted to one of the habitats.

##### Initially population is adapted to habitat L

In this biological scenario, the singular 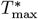 can be found by supposing the maximum stability is as in eq. (9) evaluated at 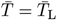 and solving eq. (S1.15), which simplifies to

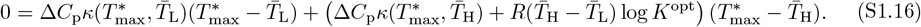

Because Δ*C*_p_ *<* 0 and 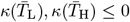, is negative everywhere except at 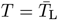 and 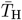 where it is 0, respectively (eq. A.30), both terms in front of 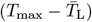 and 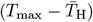 are positive. This implies that the there exists at least one solution 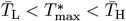 that satisfies eq. (S1.16), and that no solutions outside the interval 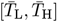 exist. The condition for the singularity to be convergent stable (eq. B.51) is

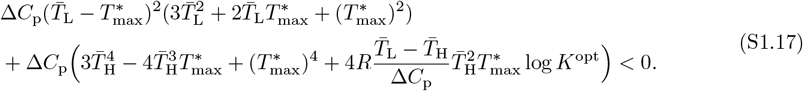

This condition is always satisfied because whenever 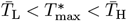, and because 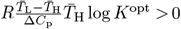. Since all singularities within the interval 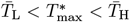 are convergence stable, there can only be one singularity (otherwise the neighboring singularities must be repellors). Our numerical exploration confirms the conclusion that if Δ*G*_max_ is initially adapted to L, then there exists a single convergent stable 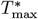, and that it satisfies 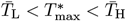.

Even if we do not know the exact explicit expression for the singularity, we can nevertheless analyse the effect of parameters on the singularity by implicit differentiation (for the method, see e.g., Otto and Day, 2011 pages 465-484, Section 12.3). In short, because the singularity must satisfy 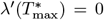 for all parameter values, differentiating this equality with respect to any parameter must also be 0. To demonstrate this method consider a parameter Δ*C*_p_ and let 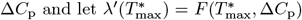. Differentiation gives

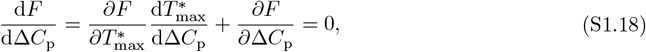

which is equivalent to

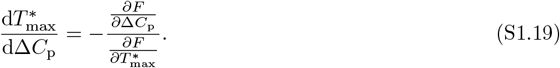

Because we want the singularity to be convergence stable, we require the denominator to be negative. And so the effect of parameter Δ*C*_p_ on the singularity 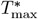 is sign equivalent to 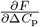, i.e.,

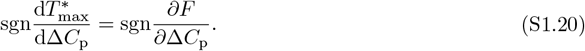

Taking the partial derivative, we get

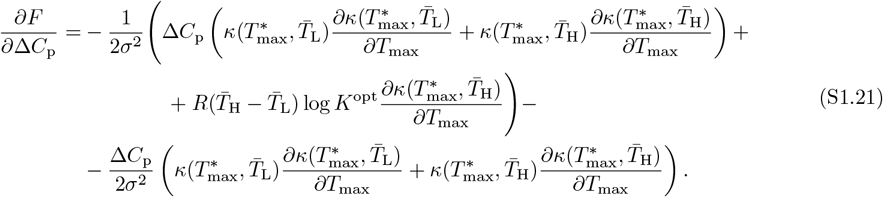

The expression inside the big brackets (first two lines) must be 0 because this is proportional to the firstorder condition for 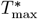 to be a singularity (eq. S1.15 being 0). Thus, the expression in the third line determines the sign of the entire expression. Because the second term inside the big brackets (on second line) is always positive (see eq. A.30), the first term in the big brackets must be always negative. Because this term is identical to the one on the third line, this means that the third line has a positive sign, and hence the entire expression is positive. This shows that increasing Δ*C*_p_ will increase the singular 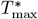.

The effect of the parameter 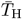 is obtained from

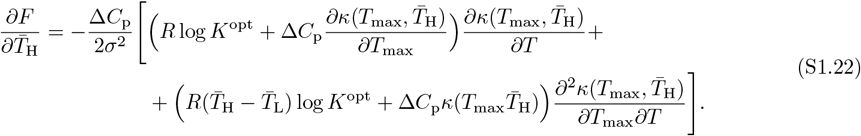

Because the expression in the brackets is positive, the entire expression is positive implying that increasing 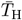 will increase the singular 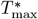. The effect of the parameter *K*^opt^ is obtained from

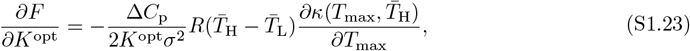

which is always positive and hence increasing *K*^opt^ will increase the singular 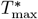. Finally, we find that parameters *σ* and *m* have no effect on the value of the singularity 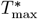.

##### Initially population is adapted to habitat H

In this biological scenario, the singular 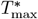 can be found by supposing the maximum stability is as in eq. (9) evaluated at 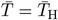 and solving eq. (S1.15), which simplifies to

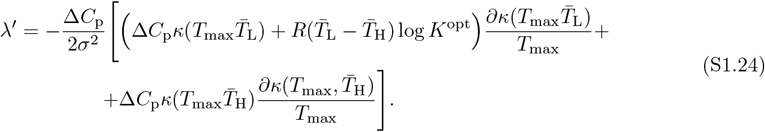

which simplifies to

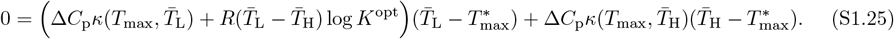

Because the first term in the brackets can be positive or negative and because 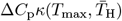 is always non-negative, solutions outside of the interval 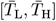 are possible. More precisely, if the term in the brackets is positive and because 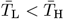, all solutions must again satisfy 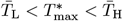. However, if the term in the brackets is negative, then either 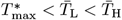, or, 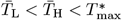. No other options are possible. In further numerical explorations, we will nevertheless constrain ourselves to solutions lying within the interval 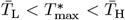, and investigate the evolutionary properties of only those that are convergent stable. To this end, let us look at the effect of Δ*C*_p_ on a convergent stable 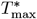 that lies within the interval 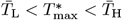. However, we can not analytically establish convergence stability in this biological scenario.

To investigate the effect of parameters on the singularity we proceed as above by implicit differentiation. We can find the effect of Δ*C*_p_ from

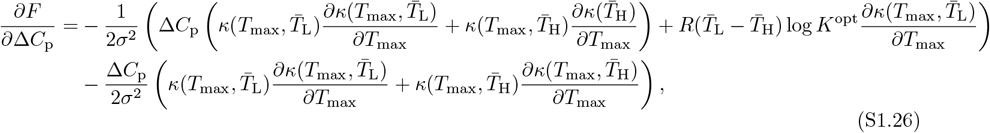

where we used eq. (S1.24). The first line must be 0 because this is proportional to the first-order condition for 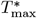 to be a singularity (eq. S1.24 being 0). Thus, the expression in the second line determines the sign of the entire expression. Because on the first line the second term in the brackets is always positive, the first term in the brackets on the first line must be always negative. Because this term is identical to the one on the second line, this means that the second line has a positive sign, and hence the entire expression is positive. This shows that increasing Δ*C*_p_ will increase the singular 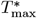.

The effect of the parameter 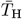 is obtained from

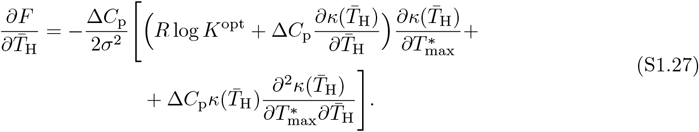

From eq. (A.30) we get that all expressions inside the brackets are positive, and hence the entire expression is positive. Increasing 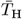 thus increases 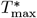. Moreover, as in the above case, the parameters *σ* and *m* have no effect on 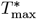.

### S1.3 Biophysical constraints in heterogeneous populations

Here we analyse the co-evolution of the thermodynamical parameters Δ*G*_max_, Δ*C*_p_ and *T*_max_ in the biological scenario where they trade off. We will analyze the following cases: (i) Δ*G*_max_ = *g*(*T*_max_) while Δ*C*_p_ is constant, (ii) Δ*C*_p_ = *g*(*T*_max_) while Δ*G*_max_ is constant, and (iii) Δ*G*_max_ = *g*(Δ*C*_p_) while *T*_max_ is constant. In the next section, we demonstrate the method by analyzing case (i); the remaining cases are analyzed similarly (Supplementary material S2). The numerical results in the main text were obtained using Mathematica version 14.0, as detailed in Supplementary material S2.

#### S1.3.1 Critical function analysis and the co-evolution maximum stability and maximum-stability temperature

In this section we analyse the (co)-evolution of Δ*G*_max_ and *T*_max_, constrained to some trade-off curve,

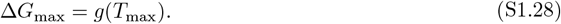

To analyze what kind of evolutionary dynamics various trade-off curves can result in, we adopt the method of Kisdi (2006, see summary on pg. 961-963) on the geometry of trade-offs in evolutionary analysis (initially developed in de Mazancourt and Dieckmann, 2004; Rueffler et al., 2004; Bowers et al., 2005 which is an extension of the method of to account for freq-dependent selection). In this method, instead of assuming a certain form of the (possibly empirically unknown) trade-off curve, one asks what kind of a trade-off curve will result in a desired evolutionary outcome. More specifically, A) the population model determines the set of all possible qualitatively different singularities: continuously stable strategy (convergent stable ESS), evolutionary branching point, invasible repellor, Garden-of-Eden. B) The population model also determines the slope (first order derivative) of the curve at a singularity. That is, given a population model, we know *a priori* what kind of singularities are possible, and if we know (or assume) the numerical value of the singularity, we also know the slope of the curve. C) The curvature (second order derivative) of the curve then determines which of the possible types of singularities is being realised. Given a population model, we can thus construct a trade-off curve *de novo* based on the desired evolutionary outcome allowed by the underlying population model.

Mutation can invade into the population whenever 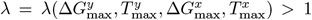, with a first-order approximation for scalar-valued parameters and small positive *δ >* 0 given as *λ*^*′*^ *>* 0 (Section B). Under the constraint, this condition reads 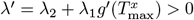 where 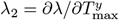 and 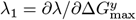. The parameters 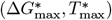 are singular whenever *λ*^*′*^ = 0, and so at a singular value the slope of the trade-off curve (eq. S1.28) is

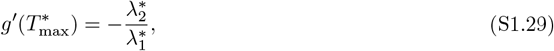

where the superscript indicates that these derivatives are evaluated at the singular value. Note that the slope of the trade-off curve (eq. S1.29) is calculated from the partial derivatives of invasion fitness, that is, without knowing the mathematical expression of the trade-off curve. The exact value of the slope is known as soon as the singularity is specified. We thus know that the trade-off curve intersects the desired singularity (by assumption), and, that its slope at the singularity is given in eq. (S1.29).

To classify the evolutionary properties of singularities, we must calculate two second order derivatives

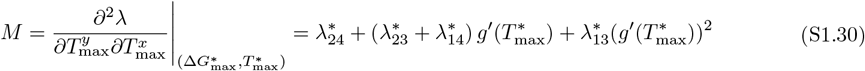

and

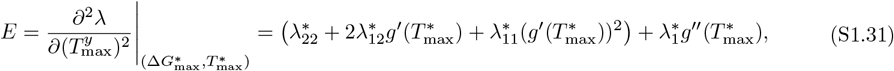

where we note that only *E* depends on the second order derivative of the trade-off function (eq. S1.28). The singularity 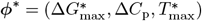 is convergent stable if *M* + *E <* 0, and evolutionary stable if *E <* 0 Geritz et al. (1998). The singularity is thus a 1) CSS whenever *M* + *E <* 0 and *E <* 0, 2) EBP whenever *M* + *E <* 0 and *E >* 0, and 3) a convergent unstable singularity (either an invasible repellopr or Garden-of-Eden) whenever *M* + *E >* 0. Importantly, note that EBP is possible only if *M <* 0 because necessarily *E >* 0. Because only *E* depends on the curvature of the trade-off curve, *M <* 0 is not only necessary but also sufficient for the model to exhibit EBP because one can always find a curvature to make *E* positive but small enough so that the conditions are satisfied.

Let us now calculate eqs. S1.28-S1.31. First, we find that the slope of the trade-off curve at singularity is

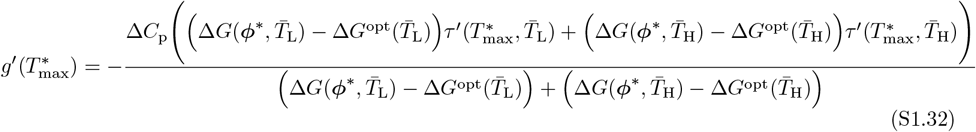

and the mixed derivative is

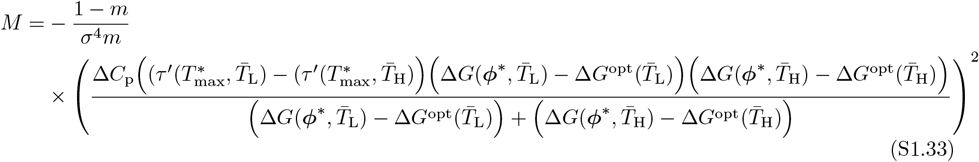

which is obtained by substituting the slope (eq. S1.32) into eq. (S1.30). We see that because 0 *< m <* 1*/*2 and 0 *< σ* the entire expression *M <* 0. We will not present the explicit analytical expression for *E* because it is lengthy.

Because *M <* 0, and we are interested in conditions that favour local adaptation via evolutionary branching, we are interested in trade-off curves that satisfy 0 *< E <* −*M*. If the first inequality is broken, the singularity is a CSS and if the second inequality is broken the singularity becomes a repellor.

In terms of the curvature of the trade-off curve, this implies that the singularity is an EBP whenever

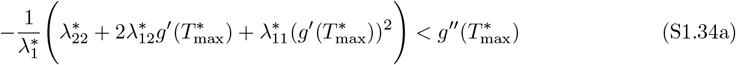

and

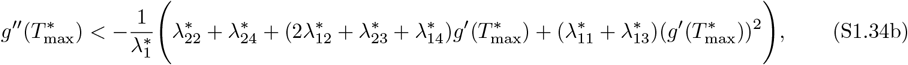

provided 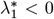, otherwise the inequality signs are reversed. If the first inequality is broken then the singularity is a CSS, and if the second inequality is broken the singularity is a repellor (REP). We have thus derived conditions for trade-off curves that facilitate local adaptation via a gradual adaptive process. The analytical expressions in terms of model parameters are lengthy and not presented here.

To facilitate numerical analysis, we construct a trade-off curve that is parametrized such that it can accommodate the various evolutionary scenarios discussed above. We choose a polynomial of degree 4 with 5 parameters (coefficients) because this allows us to specify 5 constraints that the trade-off curve must satisfy (to solve for 5 parameters we need 5 equations). For it to be valid trade-off curve, it must satisfy two constraints: intersect at a singular value, i.e., 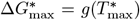, and its slope at the singularity must satisfy eq. (S1.29). The third equation specifies the curvature at a singularity 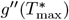, thus controlling the bifurcation between the different evolutionary scenarios according to eq. (S1.34). The first three equations thus characterize the trade-off curve locally to the singularity. In addition, as we may have information on parameter values elsewhere along the curve (but that maintain the functionality of the protein), we choose two points (Δ*G*_1_, *T*_1_) and (Δ*G*_2_, *T*_2_) that must lie on the curve, i.e., the two additional constraints are Δ*G*_1_ = *g*(*T*_1_) and Δ*G*_2_ = *g*(*T*_2_). We thus define a trade-off curve

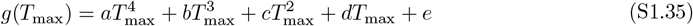

where the parameters *a, b, c, d* and *e* are solved from

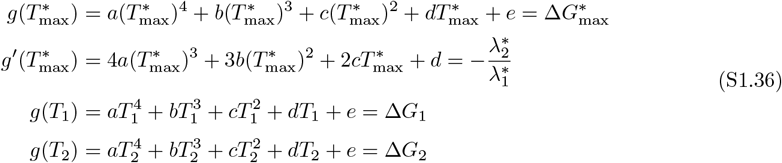

and

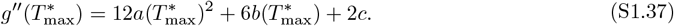

## References

Agozzino, L. and Dill, K. A. (2018). Protein evolution speed depends on its stability and abundance and on chaperone concentrations. Proceedings of the National Academy of Sciences, 115(37):9092–9097.

Alberts, B., Heald, R., Johnson, A., Morgan, D., Raff, M., Roberts, K., and Walter, P. (2022). Molecular Biology of the Cell (Seventh Edition). W. W. Norton, Incorporated.

Amarasekare, P. and Johnson, C. (2017). Evolution of thermal reaction norms in seasonally varying environments. The American Naturalist, 189(3):E31–E45.

Angilletta, M. J. (2009). Thermal adaptation: a theoretical and empirical synthesis. Oxford University Press.

Baldwin, R. L. (1986). Temperature dependence of the hydrophobic interaction in protein folding. Proceedings of the National Academy of Sciences, 83(21):8069–8072.

Barik, S. (2020). Evolution of protein structure and stability in global warming. International Journal of Molecular Sciences, 21(24):9662.

Barrick, D. (2018). Biomolecular thermodynamics: From theory to application. CRC Press.

Bastolla, U., Dehouck, Y., and Echave, J. (2017). What evolution tells us about protein physics, and protein physics tells us about evolution. Current opinion in structural biology, 42:59–66.

Becktel, W. J. and Schellman, J. A. (1987). Protein stability curves. Biopolymers: Original Research on Biomolecules, 26(11):1859–1877.

Berger, D., Stångberg, J., Baur, J., and Walters, R. J. (2021). Elevated temperature increases genome-wide selection on de novo mutations. Proceedings of the Royal Society B, 288(1944):20203094.

Bershtein, S., Serohijos, A. W., and Shakhnovich, E. I. (2017). Bridging the physical scales in evolutionary biology: from protein sequence space to fitness of organisms and populations. Current opinion in structural biology, 42:31–40.

Blaabjerg, L. M., Kassem, M. M., Good, L. L., Jonsson, N., Cagiada, M., Johansson, K. E., Boomsma, W., Stein, A., and Lindorff-Larsen, K. (2023). Rapid protein stability prediction using deep learning representations. Elife, 12:e82593.

Blanquart, F., Kaltz, O., Nuismer, S. L., and Gandon, S. (2013). A practical guide to measuring local adaptation. Ecology letters, 16(9):1195–1205.

Bomblies, K. (2020). When everything changes at once: finding a new normal after genome duplication. Proceedings of the Royal Society B, 287(1939):20202154.

Bomblies, K. (2022). The quiet evolutionary response to cellular challenges. American Journal of Botany, 109(2):189.

Bomblies, K. and Peichel, C. L. (2022). Genetics of adaptation. Proceedings of the National Academy of Sciences, 119(30):e2122152119.

Bowers, R. G., Hoyle, A., White, A., and Boots, M. (2005). The geometric theory of adaptive evolution: trade-off and invasion plots. Journal of Theoretical Biology, 233(3):363–377.

Bulmer, M. (1972). Multiple niche polymorphism. The American Naturalist, 106(948):254–257.

Bürger, R. (2000). The Mathematical Theory of Selection, Recombination, and Mutation. John Wiley and Sons, New York.

Chao, Y.-C., Merritt, M., Schaefferkoetter, D., and Evans, T. G. (2020). High-throughput quantification of protein structural change reveals potential mechanisms of temperature adaptation in mytilus mussels. BMC Evolutionary Biology, 20(1):1–18.

Chen, P. and Shakhnovich, E. I. (2010). Thermal adaptation of viruses and bacteria. Biophysical journal, 98(7):1109–1118.

Christiansen, F. B. and Feldman, M. W. (1975). Subdivided populations: a review of the one-and two-locus deterministic theory. Theoretical Population Biology, 7(1):13–38.

De Jong, M. and Saastamoinen, M. (2018). Environmental and genetic control of cold tolerance in the glanville fritillary butterfly. Journal of evolutionary biology, 31(5):636–645.

de Mazancourt, C. and Dieckmann, U. (2004). Trade-off geometries and frequency-dependent selection. The American Naturalist, 164(6):765–778.

Della Rossa, F., Dercole, F., and Landi, P. (2015). The branching bifurcation of adaptive dynamics. International Journal of Bifurcation and Chaos, 25(07):1540001.

DePristo, M. A., Weinreich, D. M., and Hartl, D. L. (2005). Missense meanderings in sequence space: a biophysical view of protein evolution. Nature Reviews Genetics, 6(9):678–687.

Dercole, F. and Rinaldi, S. (2008). Analysis of Evolutionary Processes: The Adaptive Dynamics Approach and Its Applications. Princeton University Press, Princeton, NJ.

Deutsch, C. A., Tewksbury, J. J., Huey, R. B., Sheldon, K. S., Ghalambor, C. K., Haak, D. C., and Martin, P. R. (2008). Impacts of climate warming on terrestrial ectotherms across latitude. Proceedings of the National Academy of Sciences, 105(18):6668–6672.

Dill, K., Jernigan, R. L., and Bahar, I. (2017). Protein actions: Principles and modeling. Garland Science.

Dong, Y. and Somero, G. N. (2009). Temperature adaptation of cytosolic malate dehydrogenases of limpets (genus lottia): differences in stability and function due to minor changes in sequence correlate with biogeographic and vertical distributions. Journal of Experimental Biology, 212(2):169–177.

Dong, Y.-w., Liao, M.-l., Meng, X.-l., and Somero, G. N. (2018). Structural flexibility and protein adaptation to temperature: Molecular dynamics analysis of malate dehydrogenases of marine molluscs. Proceedings of the National Academy of Sciences, 115(6):1274–1279.

Echave, J. and Wilke, C. O. (2017). Biophysical models of protein evolution: understanding the patterns of evolutionary sequence divergence. Annual review of biophysics, 46:85–103.

Feller, G. (2018). Protein folding at extreme temperatures: Current issues. In Seminars in Cell & Developmental Biology, volume 84, pages 129–137. Elsevier.

Fersht, A. (1999). Structure and mechanism in protein science: a guide to enzyme catalysis and protein folding. Macmillan.

Fields, P. A. (2001). Protein function at thermal extremes: balancing stability and flexibility. Comparative Biochemistry and Physiology Part A: Molecular & Integrative Physiology, 129(2-3):417–431.

Fields, P. A., Dong, Y., Meng, X., and Somero, G. N. (2015). Adaptations of protein structure and function to temperature: there is more than one way to ‘skin a cat’. The Journal of Experimental Biology, 218(12):1801–1811.

Galano-Frutos, J. J., Nerín-Fonz, F., and Sancho, J. (2023). Calculation of protein folding thermo-dynamics using molecular dynamics simulations. Journal of Chemical Information and Modeling, 63(24):7791–7806.

Geritz, S. A. H., Kisdi, E., Meszéna, G., and Metz, J. A. J. (1998). Evolutionarily singular strategies and the adaptive growth and branching of the evolutionary tree. Evolutionary Ecology, 12:35–57.

Ghosh, K. and Dill, K. A. (2009). Computing protein stabilities from their chain lengths. Proceedings of the National Academy of Sciences, 106(26):10649–10654.

Goldstein, R. A. (2013). Population size dependence of fitness effect distribution and substitution rate probed by biophysical model of protein thermostability. Genome biology and evolution, 5(9):1584–1593.

Graziano, G. (2008). Is there a relationship between protein thermal stability and the denaturation heat capacity change? Journal of thermal analysis and calorimetry, 93(2):429–438.

Hait, S., Mallik, S., Basu, S., and Kundu, S. (2020). Finding the generalized molecular principles of protein thermal stability. Proteins: Structure, Function, and Bioinformatics, 88(6):788–808.

Hawley, S. A. (1971). Reversible pressure-temperature denaturation of chymotrypsinogen. Biochemistry, 10(13):2436–2442.

Hochachka, P. W. and Somero, G. N. (2002). Biochemical adaptation: mechanism and process in physiological evolution. Oxford university press.

Jaenicke, R. (1991). Protein stability and molecular adaptation to extreme conditons. European Journal of Biochemistry, 202(3):715–728.

Kallioniemi, E. and Hanski, I. (2011). Interactive effects of pgi genotype and temperature on larval growth and survival in the glanville fritillary butterfly. Functional Ecology, 25(5):1032–1039.

Kawecki, T. J. and Ebert, D. (2004). Conceptual issues in local adaptation. Ecology letters, 7(12):1225–1241.

Kim, S. B., Palmer, J. C., and Debenedetti, P. G. (2016). Computational investigation of cold denaturation in the trp-cage miniprotein. Proceedings of the National Academy of Sciences, 113(32):8991–8996.

Kisdi, É. (2006). Trade-off geometries and the adaptive dynamics of two co-evolving species. Evolutionary Ecology Research, 8(6):959–973.

Leimar, O. (2005). The evolution of phenotypic polymorphism: randomized strategies versus evolutionary branching. American Naturalist, 165:669–681.

Leimar, O. (2009). Multidimensional convergence stability. Evolutionary Ecology Research, 11:191–208.

Lenormand, T. (2002). Gene flow and the limits to natural selection. Trends in ecology & evolution, 17(4):183–189.

Levins, R. (1962). Theory of fitness in a heterogeneous environment. i. the fitness set and adaptive function. The American Naturalist, 96(891):361–373.

Liao, M.-l., Somero, G. N., and Dong, Y.-w. (2019). Comparing mutagenesis and simulations as tools for identifying functionally important sequence changes for protein thermal adaptation. Proceedings of the National Academy of Sciences, 116(2):679–688.

Manhart, M. and Morozov, A. V. (2015). Protein folding and binding can emerge as evolutionary spandrels through structural coupling. Proceedings of the National Academy of Sciences, 112(6):1797–1802.

McCrary, B. S., Edmondson, S. P., and Shriver, J. W. (1996). Hyperthermophile protein folding thermo-dynamics: differential scanning calorimetry and chemical denaturation of sac7d. Journal of molecular biology, 264(4):784–805.

Meemongkolkiat, T., Allison, J., Seebacher, F., Lim, J., Chanchao, C., and Oldroyd, B. P. (2020). Thermal adaptation in the honeybee (apis mellifera) via changes to the structure of malate dehydrogenase. Journal of Experimental Biology, 223(18):jeb228239.

Metz, J. A. J., Nisbet, R. M., and Geritz, S. A. H. (1992). How should we define fitness for general ecological scenarios? Trends in Ecology and Evolution, 7:198–202.

Mylius, S. D. and Diekmann, O. (1995). On evolutionarily stable life histories, optimization and the need to be specific about density dependence. Oikos, 74:218–224.

Nagylaki, T. and Lou, Y. (2008). The dynamics of migration–selection models. In Tutorials in mathematical biosciences IV: Evolution and ecology, pages 117–170. Springer.

Oliver, T. and Palumbi, S. (2011). Do fluctuating temperature environments elevate coral thermal tolerance? Coral reefs, 30:429–440.

Otto, S. P. and Day, T. (2011). A biologist’s guide to mathematical modeling in ecology and evolution. In A Biologist’s guide to mathematical modeling in ecology and evolution. Princeton University Press.

Pancotti, C., Benevenuta, S., Birolo, G., Alberini, V., Repetto, V., Sanavia, T., Capriotti, E., and Fariselli, P. (2022). Predicting protein stability changes upon single-point mutation: a thorough comparison of the available tools on a new dataset. Briefings in Bioinformatics, 23(2):bbab555.

Priklopil, T. (2025). The role of asymmetric migration on local adaptation: an analysis of the classic two-habitat migration model. bioRxiv.

Priklopil, T. and Lehmann, L. (2020). Invasion implies substitution in ecological communities with class-structured populations. Theoretical Population Biology, 134:36–52.

Privalov, P. and Khechinashvili, N. (1974). A thermodynamic approach to the problem of stabilization of globular protein structure: a calorimetric study. Journal of molecular biology, 86(3):665–684.

Privalov, P. L. (2012). Microcalorimetry of macromolecules: The physical basis of biological structures, volume 16. John Wiley & Sons.

Pucci, F. and Rooman, M. (2014). Stability curve prediction of homologous proteins using temperature-dependent statistical potentials. PLoS computational biology, 10(7):e1003689.

Ragone, R. (2004). Phenomenological similarities between protein denaturation and small-molecule dissolution: insights into the mechanism driving the thermal resistance of globular proteins. Proteins: Structure, Function, and Bioinformatics, 54(2):323–332.

Rees, D. C. and Robertson, A. D. (2001). Some thermodynamic implications for the thermostability of proteins. Protein Science, 10(6):1187–1194.

Robertson, A. D. and Murphy, K. P. (1997). Protein structure and the energetics of protein stability. Chemical reviews, 97(5):1251–1268.

Rousset, F. (2004). Genetic Structure and Selection in Subdivided Populations. Princeton University Press, Princeton, NJ.

Rueffler, C., Van Dooren, T. J., and Metz, J. A. (2004). Adaptive walks on changing landscapes: Levins’ approach extended. Theoretical population biology, 65(2):165–178.

Sawle, L. and Ghosh, K. (2011). How do thermophilic proteins and proteomes withstand high temperature? Biophysical journal, 101(1):217–227.

Schellman, J. A. (1987). The thermodynamic stability of proteins. Annual review of biophysics and biophysical chemistry, 16(1):115–137.

Seeliger, D. and De Groot, B. L. (2010). Protein thermostability calculations using alchemical free energy simulations. Biophysical journal, 98(10):2309–2316.

Shah, P., McCandlish, D. M., and Plotkin, J. B. (2015). Contingency and entrenchment in protein evolution under purifying selection. Proceedings of the National Academy of Sciences, 112(25):E3226–E3235.

Sikosek, T. and Chan, H. S. (2014). Biophysics of protein evolution and evolutionary protein biophysics. Journal of The Royal Society Interface, 11(100):20140419.

Simon, J.-P., Potvin, C., and Blanchard, M.-H. (1983). Thermal adaptation and acclimation of higher plants at the enzyme level: kinetic properties of nad malate dehydrogenase and glutamate oxaloacetate transaminase in two genotypes of arabidopsis thaliana (brassicaceae). Oecologia, 60:143–148.

Somero, G. N., Lockwood, B. L., and Tomanek, L. (2017). Biochemical adaptation: response to environmental challenges, from life’s origins to the Anthropocene. Sinauer Associates, Incorporated Publishers Sunderland (MA).

Starr, T. N. and Thornton, J. W. (2016). Epistasis in protein evolution. Protein science, 25(7):1204–1218.

Tewksbury, J., Huey, R., and Deutsch, C. (2008). Climate warming puts the heat on tropical ectotherms. Science, 320:1296–1297.

Thurlkill, R. L., Trevino, S. R., Scholtz, J. M., and Grimsley, G. R. (2023). Determining the conformational stability of a protein from urea and thermal unfolding curves. Current protocols, 3(3):e723.

Timr, S., Madern, D., and Sterpone, F. (2020). Protein thermal stability. Progress in molecular biology and translational science, 170:239–272.

Toll-Riera, M., Olombrada, M., Castro-Giner, F., and Wagner, A. (2022). A limit on the evolutionary rescue of an antarctic bacterium from rising temperatures. Science Advances, 8(28):eabk3511.

Tuljapurkar, S. (1989). An uncertain life: demography in random environments. Theoretical population biology, 35(3):227–294.

Wang, T., He, X., Li, M., Li, Y., Bi, R., Wang, Y., Cheng, C., Shen, X., Meng, J., Zhang, H., et al. (2024). Ab initio characterization of protein molecular dynamics with ai2bmd. Nature, pages 1–9.

Williams, P. D., Pollock, D. D., and Goldstein, R. A. (2006). Functionality and the evolution of marginal stability in proteins: inferences from lattice simulations. Evolutionary Bioinformatics, 2:117693430600200013.

Yang, J., Wang, D., Liu, H., Wang, L., Jin, L., Ahola, V., Xu, C., and Wang, R. (2023). Three amino acid substitutions contributing to thermostability of phosphoglucose isomerase in the glanville fritillary butterfly. Insect science, 30(3):758–770.

Yeaman, S. and Otto, S. P. (2011). Establishment and maintenance of adaptive genetic divergence under migration, selection, and drift. Evolution, 65(7):2123–2129.

Yeritsyan, K. and Badasyan, A. (2024). Differential scanning calorimetry of proteins and the two-state model: Comparison of two formulas. Biophysica, 4(2):227–237.

Zavodszky, P., Kardos, J., Svingor, Á., and Petsko, G. A. (1998). Adjustment of conformational flexibility is a key event in the thermal adaptation of proteins. Proceedings of the National Academy of Sciences, 95(13):7406–7411.

Zheng, J., Guo, N., Huang, Y., Guo, X., and Wagner, A. (2024). High temperature delays and low temperature accelerates evolution of a new protein phenotype. Nature Communications, 15(1):2495.

Zwanzig, R. (1997). Two-state models of protein folding kinetics. Proceedings of the National Academy of Sciences, 94(1):148–150.

